# Previous malaria exposure attenuates monocyte-driven inflammation and correlates with modulation of the B cell response

**DOI:** 10.1101/2025.08.11.669804

**Authors:** Maximilian Julius Lautenbach, Pengjun Xi, Linn Kleberg, Alan-Dine Courey-Ghaouzi, Maia Serene Gower, Carolina Sousa Silva, Felicia Chammas, Anna Färnert, Christopher Sundling

## Abstract

Clinical immunity to malaria develops after repeated malaria episodes. In this process, the inflammatory response is modulated to respond less vigorously upon reinfection. Monocytes are a major source of pro-inflammatory mediators during blood-stage infection and are known to adapt to repeated pathogen exposure. Here, we investigated the impact of previous malaria exposure on monocytes during blood-stage malaria by comparing the response in previously exposed and primary infected individuals. We observed reduced levels of the proinflammatory chemokine CXCL10 in previously exposed individuals, linked to changes in CD16^+^ monocytes. Similarly, BAFF levels were lower in these individuals and associated with modulation of monocyte and dendritic cells. This affected the BAFF-BAFF-R axis, crucial for B cell responses, correlating with increasing parasite-specific antibody levels. Collectively, we present novel insights into how previous malaria exposure shapes monocyte responses during acute malaria and how these in turn correlate with modulation of the B cell compartment and humoral immune response.

## Introduction

Malaria remains a major global health concern, with estimates of 263 million cases and 597,000 deaths in 2023 according to WHO estimates (1). *Plasmodium falciparum* accounts for the vast majority of these cases and deaths. Natural immunity to malaria develops over time with repeated malaria episodes and has been historically linked to a potent antibody response against the parasite (2). Meanwhile, protection against severe malaria and immune regulation that limit immunopathology, often referred to as disease tolerance, develops more rapidly (3).

The host’s immune response is partly responsible for pathogenesis, as the parasite triggers systemic inflammation, driven by innate cells such as monocytes. Human monocytes are heterogeneous and can be classified into three major subsets: classical (CD14^+^CD16^−^), non-classical (CD14^dim^CD16^+^), and intermediate (CD14^+^CD16^+^) based on distinct surface protein markers (4). Monocytes are crucial for the initial sensing of *P. falciparum* infection through pattern recognition receptors (PRRs), such as Toll-like receptor (TLR9), which detect pathogen-associated molecular patterns (PAMPs), such as unmethylated CpG DNA (5). Through their recognition of the pathogen, they play a key role in the response to the infection, indirectly by secreting pro-inflammatory mediators to recruit and activate other immune cells and directly by anti-parasitic mechanisms of phagocytosis (6). Another effect of phagocytosis is that monocytes can present antigens to T cells, facilitating the activation of the adaptive immune response.

Monocytes play an important role in balancing protection against malaria through anti-parasite mechanisms such as phagocytosis, cytokine production, and antigen presentation, while also contributing to pathology by producing inflammatory cytokines that cause systemic inflammation and vascular dysfunction (5).

Through the secretion of inflammatory mediators, monocytes can also drive inflammation and contribute to the sequestration of infected red blood cells in organs such as the brain, placenta, and lungs by secreting cytokines that upregulate endothelial adhesion receptors (6). Especially, CD14^+^ monocytes are predominant cellular sources of cytokines and chemokines associated with severe malaria, such as tumor necrosis factor (TNF), macrophage inflammatory protein-1 alpha (MIP-1α), and MIP-1 beta (MIP-1β), and interferon-inducible protein 10 (IP-10, encoded by the CXCL10 gene (7, 8). In mouse models, the latter has also been associated with impaired parasite control during the blood stage of the infection (9).

An important aspect of keeping a balanced immune response is the monocytes’ ability to undergo epigenetic and transcriptional changes after initial exposure to malaria, leading to modulated responses upon subsequent infections, also termed innate training (5, 6). These changes can lead to two distinct and opposing states: trained immunity or tolerance. This reprogramming is induced by the engagement of various pattern recognition receptors (PRRs) with specific microbial ligands. Trained immunity is characterized by enhanced cytokine production upon restimulation, while tolerance results in decreased cytokine production (10).

Previous research has highlighted the functional modulation of monocytes in response to repeated malaria infections, including the induction of innate immune training or tolerance programs following malaria exposure (11–17). We observed differences in monocyte abundance among individuals with naturally acquired *P. falciparum* infection (18). Previously exposed individuals exhibited a higher number of CD16^+^ monocytes but experienced overall reduced systemic inflammation (18). To further understand these adaptations, we investigated longitudinal transcriptional profiles of monocytes in a cohort of returning travelers with and without prior malaria exposure. We further connected the transcriptomic data with previously generated proteomic and cell profiling data from the same individuals, followed by new mechanistic studies and serological analysis, to elucidate how monocytes during acute malaria correlate with modulation of the overall immune response, especially B cells.

## Results

### The transcriptome of monocytes is impacted by clinical *P. falciparum* infection

We investigated transcriptional changes in monocytes during acute malaria compared to healthy state 12-month control samples, by examining their gene expression profiles. We utilized a recently described targeted multi-omics single-cell dataset of peripheral blood mononuclear cells (PBMCs) in a cohort of returning travelers (19). The overall cohort was composed of returning travelers infected for the first time (primary infection), or with a previous history of malaria infection (previously exposed). The study participants presented with clinical malaria at the Karolinska University Hospital where they were included in the study. Samples were collected at treatment initiation or the day after, followed by additional sampling at 10 days, 1, 3, 6, and 12 months (**Table S1**). For the multi-omics single-cell dataset, two donors with primary infection and two donors with previous malaria exposure were analyzed.

Based on previously annotated immune cell subsets (**Figure 1A, Figure S1**), we extracted all 14,940 monocytes for in-depth analysis (**Figure 1B**) originating from 12 samples of 4 individuals with 3 time points each. In this study, “acute” refers to the sampling at diagnosis/treatment of symptomatic *P. falciparum* malaria, “D10” to the visit 10 days after treatment, and “M12” to the convalescent visit 12 months later. The multi-omics character of the data enabled us to also assess the surface protein abundance of commonly used markers for the 11,602 CD14+ monocytes and the 3,338 CD16+ monocytes (**Figure 1C**).

**Figure 1.**
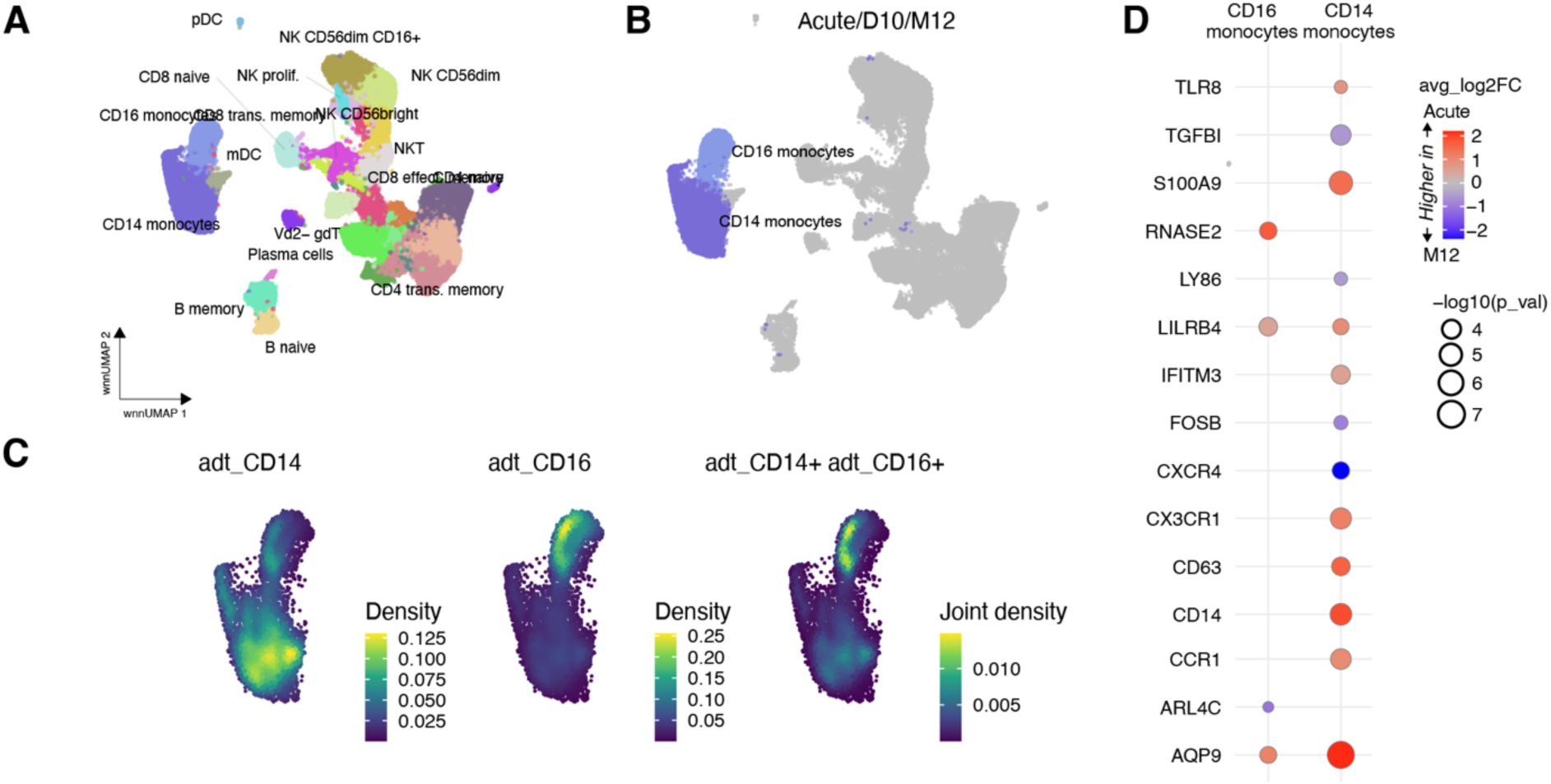
Transcriptomics profiles of monocytes during acute malaria. (**A**) UMAP representation of 73,121 CD45^+^ immune cells during and after acute malaria based on weighted nearest neighbor integrated targeted transcriptomic and epitope single-cell profiling, previously published by Lautenbach et al. (19). (**B**) Digital isolation of monocyte subsets of all three sampling time points; during acute P. falciparum malaria (Acute), approx. 10 days after treatment (D10) and at a healthy state one year after disease (M12). (**C**) Density plots for monocytes during acute malaria (only acute samples) for typical monocyte surface markers CD14 and CD16. (**D**) Differentially expressed genes (DEGs) in CD14^+^ and CD16^+^ monocytes of acute compared to the month 12 (M12) healthy state (n_Acute_ = 4, n_M12_ = 4). Red indicates higher in acute and blue higher in M12. Log transformed p-values (Wilcoxon Rank Sum test with Bonferroni correction). See also related Figure S1.

First, we assessed the differentially expressed genes (DEGs) for both monocyte subsets in the acute disease phase (Acute) compared to the healthy state after 12 months (M12). We observe more differentially expressed genes in CD14^+^ compared with CD16^+^ monocytes (**Figure 1D**). The CD14^+^ monocytes upregulated typical classical monocyte markers, such as *S100A9*, and *CD14*, while the CD16^+^ monocytes specifically upregulated *RNASE2*. Increased expression of *TLR8* in the CD14^+^ cells during the acute phase indicated a response to the infection, enabling potentially enhanced sensing of nucleic acids from parasites (20). The upregulation of *AQP9* observed in both subsets indicated the response to stimulation and activation (21). Both monocytes subset upregulates the gene *LILRB4*, an important immune checkpoint on monocytes with the ability to mediate T cell suppression (22).

### Previous malaria exposure alters the transcriptional response during acute malaria

Next, we investigated the effect of previous malaria exposure on the transcriptomic response during acute *P. falciparum* malaria. We compared the summarized gene expression (pseudobulk) of monocytes from two primary infected and two previously exposed individuals during the acute disease phase. This revealed that both CD14^+^ and CD16^+^ monocytes from primary infected individuals exhibited overall higher gene expression than previously exposed donors (**Figure 2A**). For the CD16^+^ monocytes, however, some genes such as *NKG7*, *ADGRE1*, and *VMO1* were more expressed in cells from previously exposed individuals compared to cells from primary infected individuals. Assessing differential gene expressions at the single-cell level confirmed these observations (**Figure 2B**). In CD16^+^ monocytes from previously exposed individuals, genes like *NKG7, CXCL16*, and *CD86* were significantly upregulated, consistent with higher cell expression levels compared to those with primary infection (**Figure 2B**).

**Figure 2.**
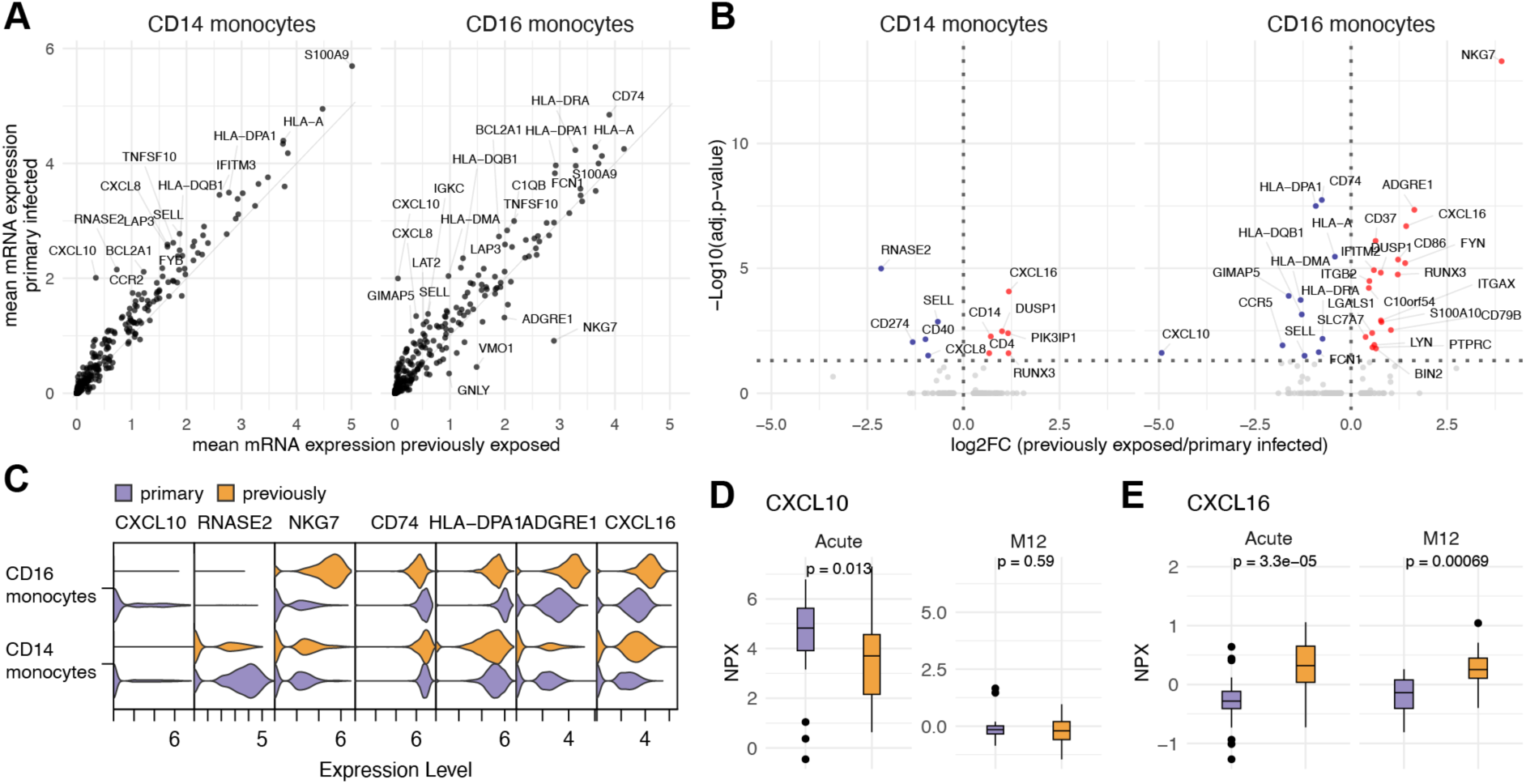
Previous malaria exposure impacts transcriptomics profiles of monocytes during acute malaria. (**A**) Gene expression in individual cells of CD14^+^ and CD16^+^ monocytes at acute malaria, from primary infected individuals (n = 2) and previously exposed individuals (n = 2). Mean expression from 3043/2583 cells (CD14^+^ monocytes) and 640/1426 (CD16^+^ monocytes). (**B**) DEGs in acute CD14^+^ and CD16^+^ monocytes in previously exposed compared to primary-infected individuals. Log transformed p-values (Wilcoxon Rank Sum test with Bonferroni correction). (**C**) Violin plots of genes are highly differentially expressed between the groups. (**D-E**) Plasma protein levels of (**D**) CXCL10 and (**E**) CXCL16 in primary infected (n=48 at acute and n=10 at 12 months) and previously exposed (n=24 at acute and n=21 at 12 months) individuals of the same cohort (19), measured with Olink Explore1536 and shown as normalized protein expression (NPX) values. Statistical analysis for D and E was done by Wilcoxon Rank Sum test with p-values indicated. See also related Figure S1.

In contrast, *RNASE2* expression was higher in CD14^+^ monocytes of primary infected individuals, while the pro-inflammatory chemokine *CXCL10* was upregulated in both monocyte subsets compared with previously exposed donors (**Figure 2C).** To investigate whether the monocyte gene expression differences we observed at the single-cell level were reflected at the systemic protein level, we repurposed a proteomic dataset generated from plasma samples from the same individuals and additional individuals from the malaria cohort of returning travelers (19). The CXCL10 levels in circulation corroborated the observation from gene expression, with previously exposed individuals having significantly (p-value = 0.013) lower systemic levels during acute malaria, when compared to primary infected individuals (**Figure 2D**). This difference was not observed when comparing the groups at their healthy state (M12) (**Figure 2D**). Further corroborating the sequencing data, we also observed that previous exposure was associated with higher CXCL16 protein levels in plasma compared with primary infected individuals (p-value = 3.3e^-05^) (**Figure 2E**). In summary, previous malaria exposure was associated with monocyte responses characterized by reduced production and secretion of the proinflammatory chemokine *CXCL10* and increased expression of *CD86* and *CXCL16* associated with T cell recruitment and activation.

### Monocyte functions are differently modulated by primary infection or previous malaria exposure

Next, we wanted to understand the functional implication of changes in the monocyte populations between primary-infected and previously exposed individuals in response to stimulation and to their microenvironment. To test this, we used infected red blood cells (iRBCs) to mimic the physiological stimuli of blood-stage malaria to assess monocyte responses. We stimulated PBMCs from primary (n = 5) and previously exposed (n = 6) individuals for 24 h using acute and convalescent (M12) samples and measured intracellular TNF and CXCL9 levels and membrane levels of lysosomal-associated membrane protein-1 (LAMP-1/CD107a) as a surrogate for overall protein secretion (for gating, see **Figure S2A**). No differences were observed for TNF or CXCL9 (**Figure S2B**) while we observed reduced CD107a levels on CD14^+^ monocytes from previously malaria-exposed individuals during clinical malaria that was normalized at convalescence (**Figure 3A**). Although the overall CD107a levels were high in primary infected individuals during acute malaria, the response was further elevated in response to iRBCs compared to stimulation with uninfected RBCs (uRBCs) (**Figure 3B**).

**Figure 3.**
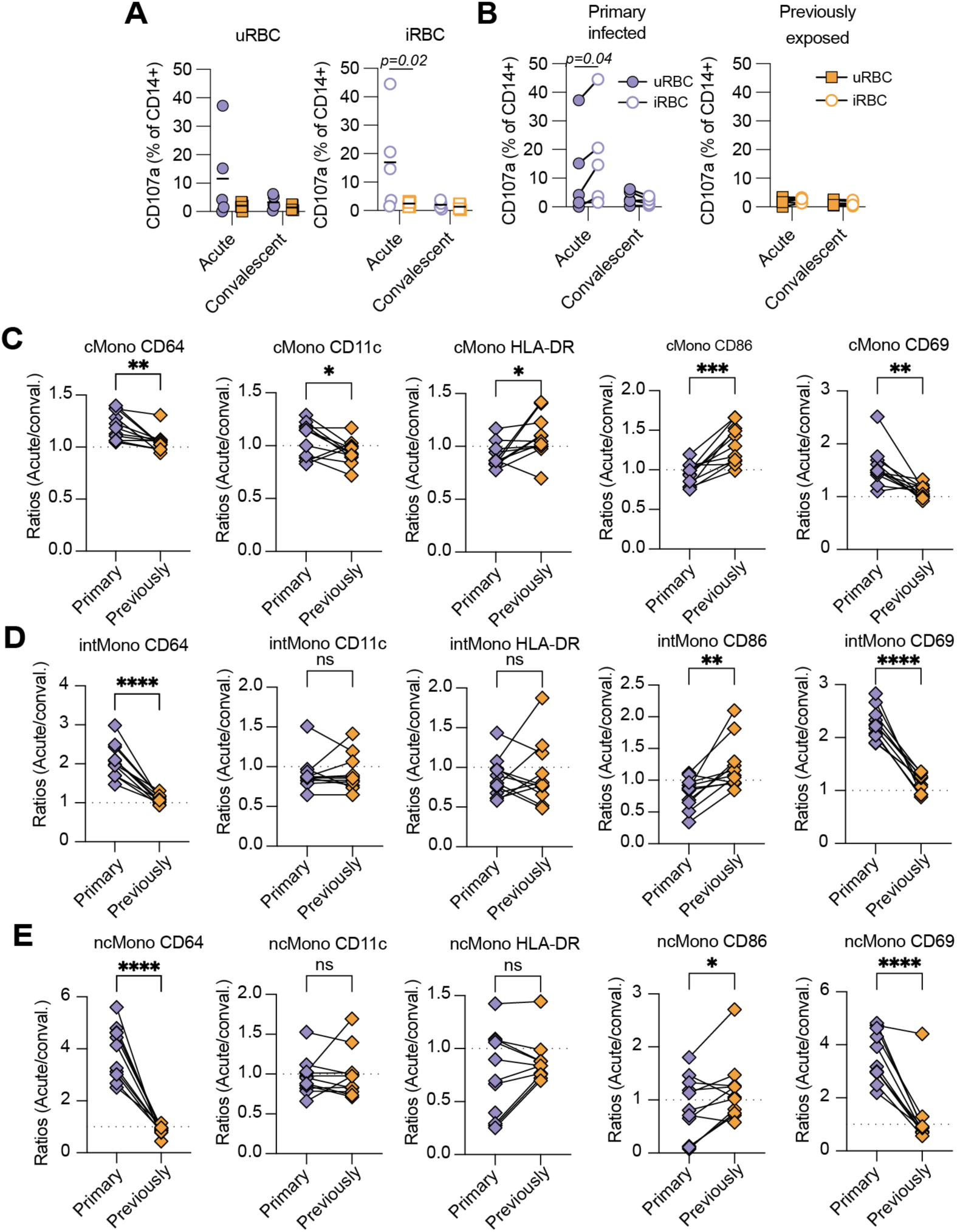
Previous malaria exposure is reflected in differences in plasma protein profiles. (**A**) CD107a^+^ monocytes after stimulation with uninfected (uRBC) or infected red blood cells (iRBC). PBMCs from primary infected (n = 5) and previously exposed individuals (n = 6) from acute and matched 12-month (convalescent) sample time points were stimulated, and degranulation was assessed via surface staining for CD107a. Primary infected individuals were compared with previously exposed individuals in (**A**) and acute samples versus convalescent samples in (**B**). (**C-E**) PBMCs isolated from healthy donors stimulated with antibody-depleted plasma pools from the acute or convalescent time-points for primary infected and previously exposed individuals. A ratio indicating geometric mean fluorescent intensity surface expression changes is shown for classical monocytes (**C**), intermediate monocytes (**D**), and non-classical monocytes (**E**). Statistical analysis in (**A**) was done using a mixed effects model matched on donor, followed by Sidak’s posttest to correct for multiple comparisons. Statistical analysis in (**B**) was calculated using two-tailed matched pair two-way ANOVA followed by Sidak’s posttest for multiple comparisons. Statistical analyses in (**C-E**) was done using two-tailed student’s t-tests with ns = p > 0.05, *p < 0.05, ** p < 0.01, *** p < 0.001, **** p < 0.0001. See also related Figure S2 and S3. Data points represent individual donors.

To test the impact of the microenvironment on monocyte function, we pooled acute or 12-month convalescent plasma from primary infected donors (n=8) and previously exposed donors (n=8) and depleted IgG antibodies to reduce antibody-mediated cytotoxic effects (23). We then cultured PBMCs isolated from healthy blood donors with 20% antibody-depleted patient plasma for 24 h and assessed levels of CD64 (FcgRI), CD11c, HLA-DR, CD86, and CD69, as indicators of functional modulation and activation (**Figure S3A**). The culture did not impact overall monocyte frequencies or cell counts between plasma pools, although there were significantly more intermediate and non-classical monocytes detected in PBMCs cultured with acute plasma from primary infected donors (**Figure S3B-C**). To assess protein surface levels, geometric mean fluorescent intensity (geoMFI) ratios were calculated between cells stimulated with acute and convalescent plasma. The ratio was then compared between cells receiving plasma from primary infected or previously exposed individuals (**Figure 3C-E**). For geoMFI values, see **Figure S3D-F**. HLA-DR and CD11c only displayed minor changes between pools in classical monocytes, with a slight increased ratio of HLA-DR and reduced ratio of CD11c in the pool from previously exposed individuals (**Figure 3C**). CD86 had a consistently higher ratio in the pool from previously exposed individuals for all monocyte subsets, although geoMFI changes were mainly observed for classical monocytes, while CD64 and CD69 had a consistently higher ratio and geoMFI in the pool from primary infected individuals (**Figure 3C-E**).

In summary, monocytes from previously malaria exposed individuals displayed reduced baseline activation and response to stimulation, while the plasma stimulated increased levels of the co-stimulatory molecule CD86. In contrast, monocytes from primary infected individuals displayed high baseline activation and exocytosis and a plasma profile promoting monocyte activation, indicated by increased surface levels of CD64 and CD69.

### Previous malaria alters circulating protein profiles during acute malaria

Due to the reduced monocyte responses in previously exposed donors upon ex vivo iRBC stimulation and different impact of plasma pools from primary infected and previously exposed donors on cellular responses, we thought to further investigate differences in protein profiles associated with previous malaria exposure. For a more accurate analysis, we focused on individuals with available protein profiling data at both the acute and 12-month time points. Between primary infected (n = 10) and previously exposed (n = 21) individuals, there were no differences in age (p-value = 0.5), sex (p-value = 0.4), or parasitemia (% iRBCs; p-value = 0.4) (**Table S1**). We identified 40 differentially abundant proteins (DAPs) between the groups at acute, compared to their 12-month follow-up, using a linear mixed-effect model approach to account for individual subject effects (**Figure 4A-B**).

**Figure 4.**
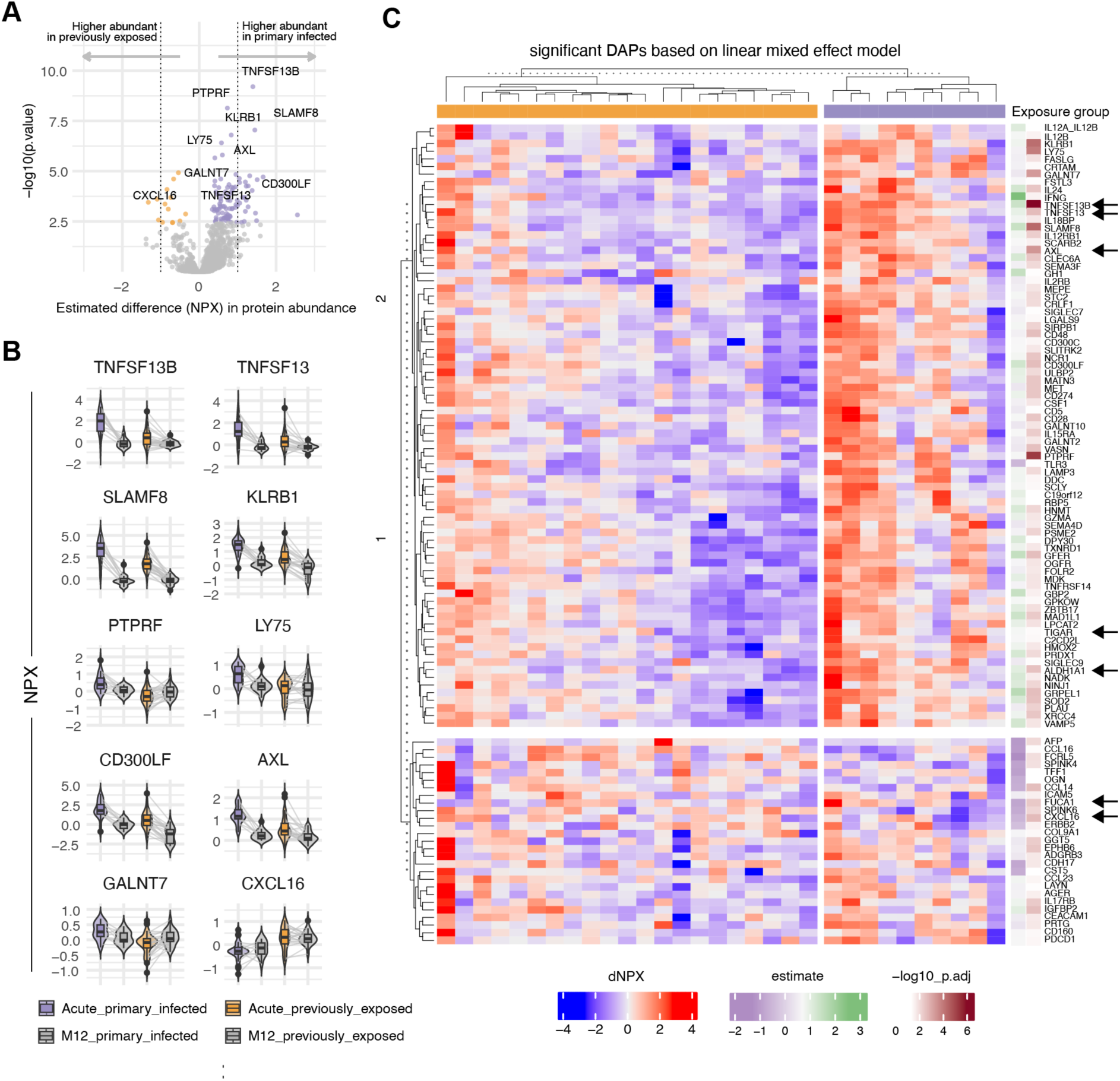
Plasma protein differences associated with malaria in individuals with primary infection versus previous exposure. Differential protein abundance between primary infected (n = 10) and previously exposed individuals (n = 21), based on a linear mixed-effects model accounting for inter-subject variation, with FDR correction (Benjamini–Hochberg, 0.05) applied throughout. **(A)** Volcano plot comparing NPX values between primary infected and previously exposed donors (controlling for donor and time) and highlighting the top 10 differentially abundant proteins (DAPs) ranked by adjusted p-value and NPX difference. **(B)** Protein abundance of the top 10 significant proteins (all have adjusted p ≤ 0.05) across acute and convalescent (M12) states. **(C)** Heatmap of ΔNPX (Acute–M12) for all significant DAPs, annotated with NPX differences and −log10(adjusted p-values). Parameter estimate indicates the estimated difference in protein abundance.

Overall, we observed that most proteins (29 of 40) were higher in primary infected individuals, including cytokines (TNFSF13B, TNFSF13, FASLG, CSF1), enzymes (ASAH2, GALNT10, GZMA), and receptors (PTPRF, CRLF1, CD27) (**Figure 4C**). In contrast, only 11 proteins were higher in previously exposed individuals, including chemokines (CXCL16, CCL14), receptors (FCRL5), enzymes (FUCA1), and enzyme inhibitors (SPINK4, SPINK6).

Among the 40 proteins, seven (TNFSF13B, CXCL16, FUCA1, ALDH1A1, TNFSF13, AXL, and TIGAR) have been linked to monocytes and DCs via blood gene expression profiles generated by the Human Protein Atlas (proteinatlas.org) and are indicated by arrows in **Figure 4C**. Among them, TNFSF13B or B cell activating factor (BAFF), showed the greatest difference between primary-infected and previously exposed individuals with 4-fold higher abundance in primary-infected compared to previously exposed individuals.

### Association between monocyte and DC-produced BAFF and BAFF-R levels on B cells

The significant differences in circulating TNFSF13B (BAFF) levels associated with previous malaria exposure prompted an investigation into potential immune cell sources. To approximate this, we analyzed gene expression levels from the scRNAseq dataset in various immune cell subsets. Comparing the average gene expression per immune cell subset from all four donors with acute malaria, we found that myeloid cells (CD14^+^ and CD16^+^ monocytes, mDCs, and pDCs) exhibited the highest average expression among PBMCs (**Figure S4A**), corroborating existing literature^21^. The gene expression of *TNFSF13B* in mDCs, CD14^+^, and CD16^+^ monocytes showed a decreasing trend across the sampling time points (Acute, 10 days, 12 months), consistent with higher expression during acute infection (**Figure S4B-C**). To further illustrate this temporal pattern, we compared the average gene expression levels between acute and 12-month samples, which showed higher average *TNFSF13B* expression across samples during the acute phase, most prominently in mDCs with a 0.48-fold increase, followed by CD16^+^ monocytes a 0.37-fold increase, and CD14^+^ monocytes a 0.35-fold increase (**Figure S4D**).

BAFF is the sole ligand for the BAFF-receptor (BAFF-R), encoded by the *TNFRSF13C* gene, and serves as a critical pro-survival receptor for B cells (24). The *TNFRSF13C* gene expression was low for B cells in our cohort, while the surface protein expression was high for naïve and memory B cells, but low for plasma cells (**Figure S5A-C**). To validate these observations, we downloaded and repurposed a previously published malaria single-cell RNAseq dataset (25) including samples from individuals with acute malaria (day 0), and days 7 and 28 after treatment, as well as from non-infected controls **(Figure S5D).** We observed relatively even BAFF-R expression levels over time and compared with non-infected controls. Consistent with our own dataset, *TNFRSF13C* expression was low in plasma cells compared with other B cells.

To further explore the potential association between BAFF and the BAFF-R in clinical malaria, we reanalyzed previously generated flow cytometry data from 50 individuals of the same traveler cohort (26), where we examined BAFF-R expression on B cell subsets from acute infection through to 12 months post-infection (see **Figure S6A** for the gating strategy). During acute malaria, BAFF-R levels on circulating B cells were significantly depleted, with levels returning to those of healthy controls (n = 14) approximately 3-6 months after infection (**Figure 5A-B**). This pattern was similar across B cell subsets (**Figure S6B-C**). When comparing BAFF-R levels on B cells from primary (n = 17) and previously exposed (n = 33) individuals during acute infection, we observed significantly lower levels in the primary infected group (p = 0.0068) (**Figure 5C**). This was also reflected in the surface expression of single-cell data (**Figure S5C**).

**Figure 5.**
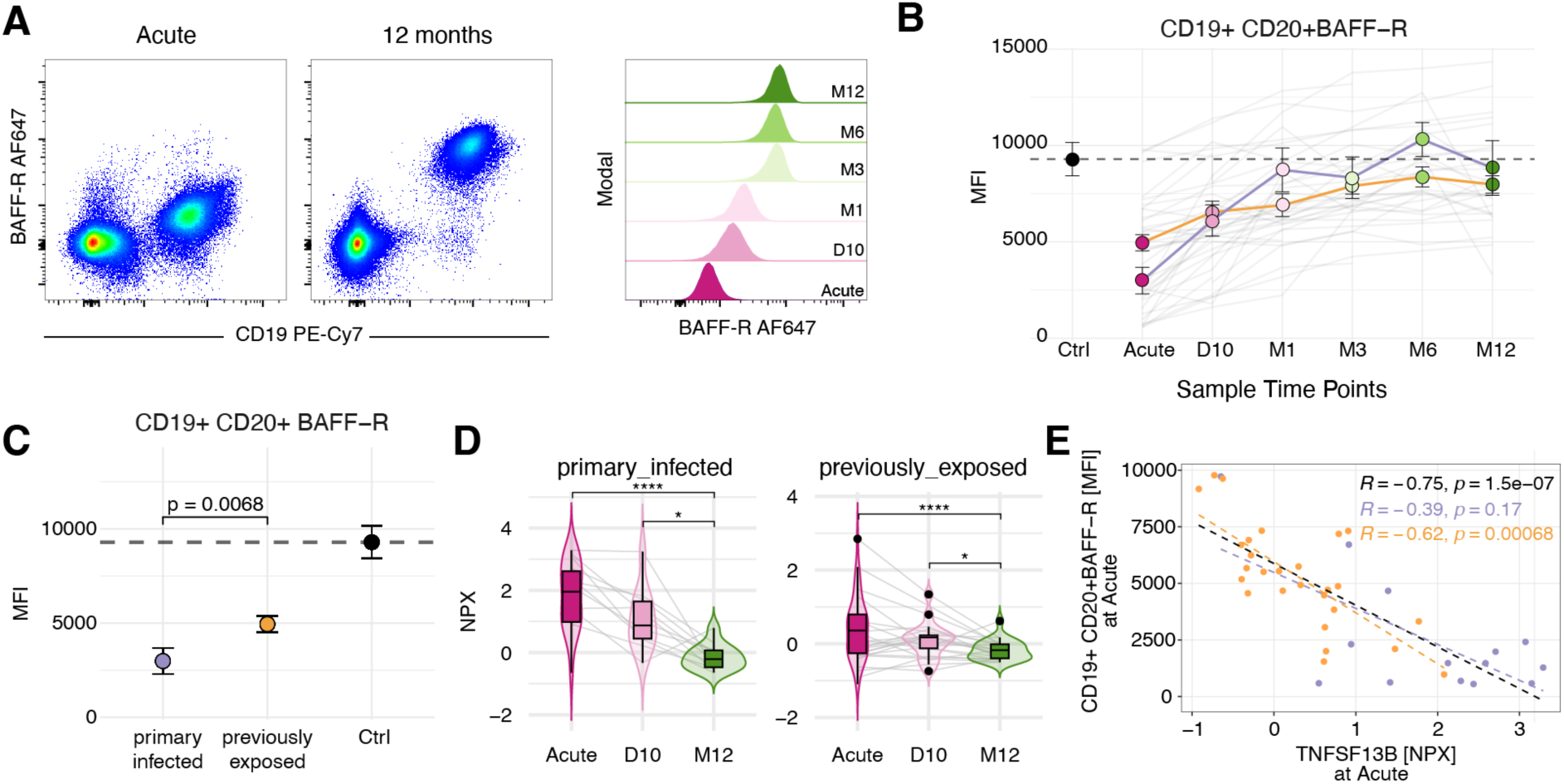
BAFF-R levels on B cells are reduced during acute malaria and highly negatively correlated with circulating BAFF levels. (**A**) Representative FACS plots of BAFF-R expression on total live lymphocytes at acute P. falciparum malaria or after 12 months (left and middle panel) and for CD19^+^ CD20^+^ B cells at all time-points (right panel). (**B**) Longitudinal assessment of BAFF-R expression on CD19^+^ CD20^+^ B cells at and after acute P. falciparum malaria. The points denote the mean with error bars showing the standard error (SE). The grey lines indicate data from individual donors. Healthy individuals (Ctrl) as a reference with dotted line of their mean (n = 14) (**C**), Comparison of CD19^+^ CD20^+^ BAFF-R^+^ B cells during acute malaria for primary infected (purple, n = 15) and previously exposed individuals (orange, n = 28). Mean of control group as dotted line as in (**B**). (**D**) Relative protein abundance NPX of BAFF in primary infected and previously exposed individuals over the sampling time points, assessed using linear mixed-effects models accounting for repeated measures. *P < 0.05, **P < 0.01, ***P < 0.001, ****P < 0.0001.(**E**) Scatterplot of CD19^+^ CD20^+^ BAFF-R^+^ at acute malaria and the corresponding BAFF levels, annotated with Spearman correlation. Data points are colored based on primary-infected (purple, n = 14), and previously exposed (orange, n = 27) with corresponding trendlines. The black trendline, correlation coefficient and p-values corresponds to all data. See also related Figure S6. Data points represent individual donors.

BAFF protein level dynamics in plasma over the three sampling time points (Acute, 10 days, and 12 months) for primary infected and previously exposed individuals confirmed that primary infected individuals exhibited highly elevated levels in response to malaria. In contrast, previously exposed individuals displayed only slightly elevated levels during acute malaria compared with at 12 months (**Figure 5D**). To investigate if the circulating BAFF levels were associated with the BAFF-R levels on B cells, we performed Spearman correlation analysis at the acute infection between matched plasma BAFF levels and mature B cell BAFF-R levels. The analysis revealed a significant inverse correlation (merged data: rho = −0.75 p-value < 0.001, previously exposed: rho = −0.62 p-value = 0.00068, primary infected: rho = −0.39 p-value = 0.17) (**Figure 5E**), indicating a potential link between high plasma BAFF levels and reduced BAFF-R expression on B cells. These observations are consistent with previous findings from a controlled human malaria infection study, where primary malaria was associated with increased plasma BAFF levels and an inverse correlation between plasma BAFF and B cell BAFF-R expression (27).

### BAFF-R downregulation is mediated by soluble plasma factors

The better understand which factors that could modulate BAFF-R levels on B cells, we used in vitro experimental approaches. We first assessed if soluble factors within the patient plasma could modulate BAFF-R levels. This was done by culturing healthy PBMCs with plasma pools from primary infected or previously exposed patients at acute malaria or convalescence for 24 h followed by gating for B cells (**Figure 6A**).

**Figure 6.**
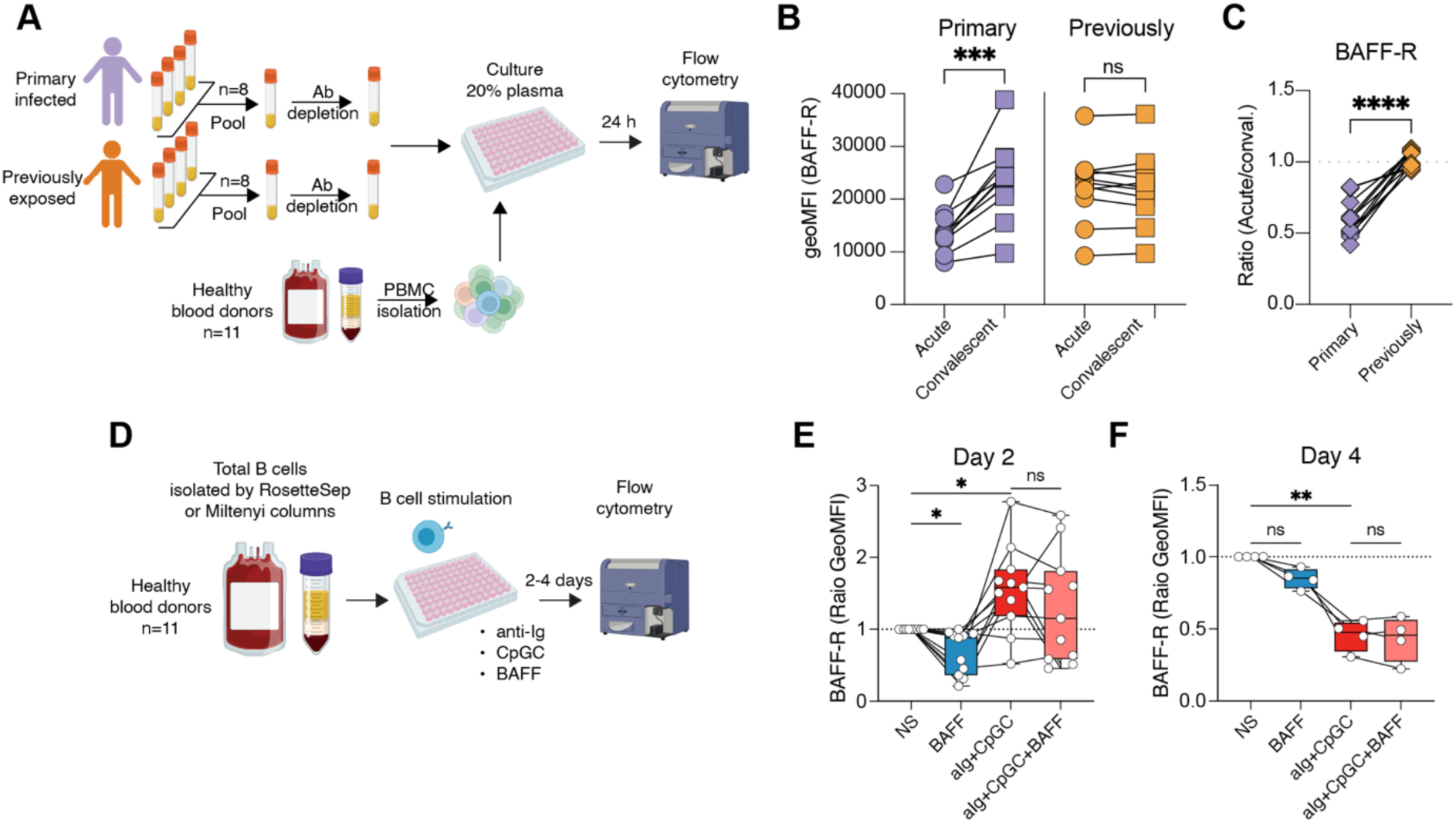
BAFF-R levels are influenced by BAFF, but not via transcriptional regulation or receptor shedding. (**A**) Schematic illustration of the in vitro experimental setup for plasma stimulations of PBMCs. Plasma was pooled from primary infected (n=8) or previously exposed (n=8) donors while PBMCs were isolated from healthy blood donors (n=11). (**B**) Geometric mean fluorescent intensity (geoMFI) of BAFF-R levels on gated B cells 24 h after culture with indicate plasma pools. (**C**) Comparison of the BAFF-R geoMFI ratio for acute over 12-month plasma for primary infected and previously exposed donors. Statical analysis in B was done using a paired two-tailed student’s t-tests and in C an unpaired two-tailed student’s t-test. Ns = p > 0.05, ***p < 0.001, ****p < 0.0001. (**D**) Schematic illustration of the experimental setup for recombinant BAFF stimulations of purified B cells. (**E**) BAFF-R geometric mean fluorescent intensity (GeoMFI) ratios between non-stimulated and BAFF, anti-Ig+CpGC, or anti-Ig+CpGC+BAFF stimulated sorted B cells after (**E**) 2 days (n=11) or (**F**) 4 days (n=4) of culture. Statistical analyzes in E and F were calculated using repeated-measures one-way ANOVA followed by Sidak’s posttest with ns = p > 0.05, *p < 0.05, **p < 0.01 See also related Figure S7. Data points represent individual donors.

B cells from PBMCs stimulated with plasma were gated as indicated in **Figure S3** with an additional gate for BAFF-R+ cells (**Figure S7A**). The stimulation had no impact on overall B cell numbers or BAFF-R+ cells (**Figure S7B-C**). In addition to BAFF-R levels, we also assessed surface expression of HLA-DR, CD11c, FcRL5, CD69, and CD86. Of these, only plasma from primary infected individuals at the acute time-point could promote significant changes in B cell surface expression. FcRL5 displayed a modest upregulation compared with convalescent plasma, although the geoMFI levels remained similar to those from previously exposed plasma (**Figure S7D-E**). BAFF-R levels, however, were robustly reduced in both geoMFI and in ratio compared with plasma from previously exposed donors (**Figure 6B-C**).

Since the plasma pools from primary infected individuals contained high levels of BAFF at the acute time-point compared to previously exposed donors and convalescent time-points (**Figure S7F**), we next assessed if recombinant BAFF could be the driver of BAFF-R downregulation. We sorted total B cells from healthy individuals and stimulated them with BAFF alone, or in combination with anti-Ig (BCR ligand) and CpGC (TLR9 ligand), to mimic B cell activation during malaria infection (**Figure 6D**). After 2-4 days of culture, we compared BAFF-R levels between non-stimulated and stimulated cells. We observed that B cells stimulated with BAFF exhibited significant downregulation of BAFF-R (p-value < 0.05) compared to non-stimulated cells, while B cells stimulated with BCR/TLR9 ligands initially (day 2) showed significant upregulation of BAFF-R (p-value < 0.05) compared to non-stimulated cells (**Figure 6E**), but then reduced overall BAFF-R levels at day 4 of culture (gated on non-plasmablasts) (**Figure 6F**). B cells stimulated with both BCR/TLR9 ligands and BAFF showed no significant difference in BAFF-R levels (p-value > 0.05) compared to cells stimulated with BCR/TLR9 ligands alone (**Figure 6E**). As expected from our own and previous data (28), we observed low BAFF-R levels on plasmablasts on day 4 of stimulation **(Figure S7G**). Further analysis of the BAFF-R levels together with other B cell markers on day 2 and 4 of stimulation indicated that BAFF-R low cells also displayed reduced levels of CD19 but normal levels of CD20 and no increased staining for a cell death marker (**Figure S7H-J**). This phenotype was not observed in B cells from malaria patient samples (**Figure S7I**). Such a population was also seen when cultured with patient plasma, although the reduced BAFF-R levels observed were gated from the BAFF-R+ CD19 high population and not CD19 low cells (**Figure S7J**).

In summary, the plasma from acute malaria in primary infected individuals, but not previously exposed donors, could drive BAFF-R downregulation. This effect could not be completely replicated through the addition of recombinant BAFF to B cells, indicating that additional factors or different oligomeric forms of BAFF in the plasma could be important in mediating the BAFF-R downregulation.

### High levels of BAFF in circulation are correlated with reduced parasite-specific antibody responses

To investigate the association between circulating BAFF and humoral immunity during acute malaria, we correlated plasma BAFF levels with malaria-specific antibody responses in individuals with primary or previous malaria exposure. Antibody responses were summarized using a cumulative response score (CRS) across 5 *P. falciparum* merozoite antigens (AMA1, MSP1-19, MSP2, MSP3, and GAMA). We observed that BAFF levels negatively correlated with CRS for IgG (rho = –0.63, p-value < 0.0001), IgG1 (rho = –0.62, p-value < 0.0001), IgG3 (rho = –0.69, p-value < 0.001), and for IgM (rho = –0.5, p-value = 0.00013) (**Figure 7**). These correlations remained significant for IgG3, but not for total IgG, IgG1, or IgM, when stratifying by malaria exposure status, although the trend lines remained similar (**Figure 7**). We next examined whether parasitemia could be a driving factor to explain either the BAFF (**Figure S8A**) or antibody levels (**Figure S8B**). However, no significant correlations were observed in either case, indicating that the BAFF-antibody correlation was not driven by parasitemia.

**Figure 7.**
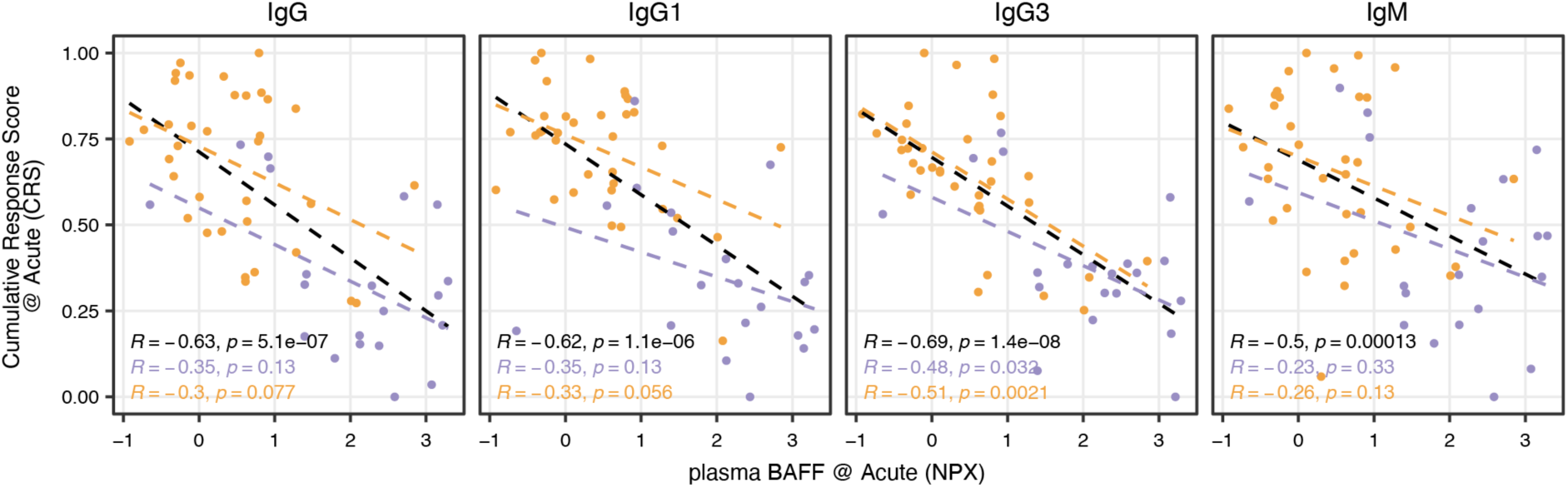
Circulating BAFF levels are inversely correlated with malaria-specific IgG responses. Correlation between acute BAFF levels and Cumulative Response Score (CRS) of total IgG, IgG1, IgG3, and IgM responses to 5 merozoite antigens in matching individuals (total n=55, with primary infected n=20 and previously exposed n=35). Correlation was assessed using Spearman correlation with samples colored based on primary infected (purple) and previously exposed (orange) individuals, with coefficients and p-values calculated for the combined cohort (black) and for primary infected (purple) and previously exposed (orange) separately. See also related Figure S8. Data points represent individual donors.

Together, these findings suggest that elevated BAFF during acute infection may be associated with less effective antibody responses in individuals experiencing their first malaria exposure.

## Discussion

Monocytes play a crucial role in balancing the anti-parasite response and contributing to systemic inflammation and pathology in malaria(5). While disease-protecting immunity develops through a broad antibody repertoire, protection from severe disease arises more rapidly, likely through a combination of increasing humoral responses and immune regulatory changes that limit immunopathology (18). Here, we analyzed immune response data from individuals with acute *P. falciparum* malaria, focusing on the impact of previous malaria exposure. We report transcriptional changes in monocytes that influence the acute inflammatory response and immunomodulatory mechanisms that could have an effect on the B cell response during malaria.

We reported previously that individuals with prior exposure to malaria were better able to control parasitemia through the action of cytophilic (IgG1 and IgG3) antibodies in conjunction with CD16^+^ monocytes, contributing to less systemic inflammation (18). The transcriptional changes in monocytes observed here support these findings. Higher expression of CXCL16, a membrane-bound chemokine, functions as a scavenger receptor on macrophages, binding phosphatidylserine (PS) and oxidized lipoprotein (29). The exposure of phosphatidylserine on the plasma membrane of cells is a known signal of apoptosis, enabling phagocytes to recognize dying cells (30). Infected erythrocytes are reportedly also able to expose PS on their outer surface (31). Enhanced recognition and clearance of apoptotic cells via membrane-bound CXCL16 may help remove damage-associated molecular patterns (DAMPs) and endogenous factors that could trigger more inflammation. Conversely, higher plasma levels of soluble CXCL16, a chemoattractant for activated CXCR6^+^ T cells, may result from increased membrane-bound CXCL16 on monocytes, shed through ADAM10-mediated cleavage (32).

Transcriptomic profiles of monocytes indicated reduced *CXCL10* expression in previously exposed donors compared with primary-infected individuals. CXCL10, a pro-inflammatory chemokine, recruits CXCR3^+^ leukocytes, contributing to inflammation and tissue damage (33). Monocytes from previously exposed individuals showed almost no *CXCL10* expression, correlating with significantly lower plasma levels. CXCL10 interacts with CXCR3, which is expressed by atypical B cells (26, 34, 35), plasmacytoid dendritic cells (pDCs) and activated effector CD4^+^ T cells during acute malaria (19). Reduced CXCL10 availability may therefore decrease the activation and recruitment of these immune cells, reducing overall immune activation, while it amplifies monocyte inflammatory responses (33), potentially causing more host damage in primary infected individuals. The production and secretion of CXCL10 is IFNγ-dependent (8), suggesting that the observed lower levels of IFNγ in previously exposed individuals potentially explain the reduced CXCL10 expression and secretion (18). IFNγ has also been suggested to modulate myeloid cell training and tolerance responses (36), potentially explaining the lower monocyte degranulation we observed following ex-vivo iRBC stimulation. Disease tolerance in malaria has also been associated with metabolic changes (37). This was potentially also reflected in our data, where previously exposed individuals had reduced plasma levels of TIGAR (TP53-induced glycolysis and apoptosis regulator), which increases during cell stress to reduce glycolysis and increase available NADPH for DNA repair (38).

Although not investigated here, epigenetic changes in monocytes have been reported to be involved in immunomodulation and innate-driven tolerance (39). Epigenetic reprogramming of human monocytes from individuals exposed to repeated *P. falciparum* infections can shift the monocytes to become more regulatory (11). These changes contribute to attenuated inflammatory responses during subsequent malaria exposures, reducing the risk of severe inflammatory symptoms. These observations should also be interpreted in the context of early-life and environmental exposures, which may contribute to long-lasting immune conditioning of monocytes independently of malaria (40). One such factor could be cytomegalovirus (CMV) infection which was proportionally higher in previously exposed donors in this study. Monocytes and stem cell progenitors have been described as reservoirs for latent CMV infection (41, 42), and activation of such in vitro infected cells showed less responsiveness to interferon gamma signaling (43).

Guha *et al.* showed that ex-vivo stimulated monocytes of Mali adults secreted less TNF, IL-1β, and IL-6, and more IL-10 compared to unexposed US adults (11). In our cohort, we observed reduced plasma levels of TNF and IL-6 in previously exposed individuals (although not significant) but no elevated levels of IL-10 compared to primary infected individuals (18).

Other cells, rather than monocytes, might be responsible for high blood IL-10 levels. Transcriptomic profiles suggest that regulatory T (Tr1) cells could be an important source of IL-10 during acute malaria (25). CD16^+^ monocytes express high TNF levels, potentially contributing to plasma TNF (19). In contrast, we did not observe strong IL-6 expression in monocytes during acute malaria; instead, IL-6 was primarily expressed by B cells (19). Endothelial cells have also been reported to contribute to circulating IL-6 (44), although this was not analyzed here. Collectively, our findings suggest that monocyte changes, through direct and indirect effects, contribute to dampened systemic inflammation linked to previous malaria exposure. This response modulation benefits disease protection by decreasing potential host damage and symptoms and may contribute to the development of a better humoral response.

One protein that was strongly influenced by previous malaria exposure was TNFSF13B (BAFF), which was significantly more abundant in primary infected individuals. The myeloid cells upregulated *TNFSF13B* expression during malaria, likely contributing to the higher circulating levels as they have been described to be the major sources of BAFF (45). BAFF is critical for immature B cell maturation and mature B cell maintenance but is also associated with plasma cell differentiation and survival (24, 28, 45). IFNγ can upregulate BAFF expression in monocytes (46), suggesting that the elevated BAFF levels might result from the significantly higher IFNγ levels observed in primary infected individuals. High IFNγ and other pro-inflammatory mediators in primary infected individuals could represent a disadvantageous inflammatory response, impairing the induction of a long-lasting B-cell response. This is supported by observations in other studies where individuals with a high antibody response had lower plasma BAFF, possibly due to reduced IFNγ levels (47).

BAFF can bind to three receptors, BAFF-R, TACI, and BCMA. TACI and BCMA also bind the ligand APRIL, while BAFF-R is specific for BAFF (48–50). BAFF-R is highly expressed on immature and mature B cells, where it is needed for survival and maintenance (51). As B cells become activated and differentiate to plasma cells they downregulate BAFF-R and upregulate BCMA and TACI and become more dependent on APRIL for survival (52). BAFF-R levels can be regulated through shedding via cleavage with ADAM10 and ADAM17, as reviewed by Meinl et al. (53). The cleavage is induced via binding by BAFF trimer or 60-mer to BAFF-R. The effect can also be induced in vitro via TLR9-ligation and is then dependent on ADAM10 (54). In this study, we observed BAFF-R downregulation on B cells from malaria patients. We could recapitulate this effect upon culture of healthy PBMCs with plasma from primary infected donors. We also observed BAFF-R downregulation in response to BCR and TLR9 stimulation, although those cells also downregulated CD19, contrasting with the ex vivo patient B cells. This could potentially be explained by CD19 internalization, which has been described as a mechanism to regulate B cell activation upon activation. In vivo the levels of CD19 then restore via de novo production upon cessation of the stimuli (55).

In the context of malaria, impairment of the BAFF-BAFF-R axis has been linked to disease severity and poor malaria antigen-specific B cell differentiation (56). Similar to our study, Nduati *et al.* observed reduced BAFF-R levels on peripheral B cells (56). However, children with higher BAFF-R levels on their B cells maintained parasite-specific IgG over four months, indicating more long-lasting B cell responses in individuals with reduced inflammatory responses. The study concluded that the observation was due to dysregulation of BAFF-R on B cells, in favor of short-lived antibody responses (56). Here, we could also show a negative association between antibody levels in adults during the acute inflammatory response, where high BAFF levels were associated with lower levels of parasite-specific IgG and IgM antibodies. Although individuals with previous malaria exposure could retain memory B cells, contributing to more rapid antibody production, IgG3 and IgM antibody levels are less associated with pre-existing long-lived memory. The negative correlation of these with BAFF levels therefore more directly indicates a possible negative effect of the inflammatory response on antibody production.

However, it remains possible that rapid extrafollicular IgM and IgG3 antibody production in naïve individuals and memory-derived high-affinity IgG1 antibodies could be important drivers in modulating the early monocyte and systemic inflammatory response. Acute malaria has been associated with rapid and extensive expansion of atypical B cells (26), which have been suggested to be derived from extrafollicular reactions (57). In malaria, atypical B cells are enriched for unswitched B cell receptors (58). They also have high plasma cell differentiation potential in the early stages of atypical B cell differentiation (58). However, they have also been described to require inflammatory signals to be generated (59), making it difficult to determine what comes first and what role each component contributes with to the overall modulation of the host response, leaving the temporal and mechanistic relationships between these factors to be addressed by future studies.

Although BAFF levels did not correlate with peripheral parasitemia in our cohort, this does not necessarily contradict previous findings showing an association between BAFF-R expression in splenic B cells and parasitemia in malaria patients (60). Differences in tissue compartment analyzed, time of sampling and disease context may contribute to these divergent observations. Together with prior CHMI study (27), these findings suggest that elevated BAFF and reduced BAFF-R expression are common features of acute malaria, while their relationship to parasitemia may depend on anatomical compartment and disease context.

In conclusion, our findings highlight the central role of monocyte-mediated immune modulation in developing tolerance and humoral immunity to malaria. The observed changes in transcriptional and cytokine profiles suggest that prior exposure to malaria can have an imprinting effect on monocytes to balance effective parasite control with reduced systemic inflammation. Implementing these insights could enhance therapeutic strategies aimed at modulating the immune response to prevent severe malaria.

### Limitations of the study

A key limitation of this study is the influence of host genetic variability on immune responses, which is difficult to control for in prospective study designs. Furthermore, although single-cell transcriptomic profiling was performed on a large number of cells, the data were derived from only four individuals (two per exposure group), potentially limiting generalizability. These individuals were selected to avoid extremes based on prior data, but differential gene expression may not reflect broader cohort trends. We could confirm overlapping phenotypic changes using an experimental model but validation in larger, independent cohorts is warranted. The association between circulating BAFF levels and antibody responses is based on correlations. Conclusively determining if BAFF controls antibody secretion or vice versa therefore necessitates further functional experiments to investigate mechanistic connections.

## Methods

### Resource availability

Materials availability

This study did not generate new unique reagents.

Data and code availability

Data that was repurposed from previously generated/published datasets

- Single-cell multi-omics data (Rhapsody target Immune response panel and AbSeq) and plasma protein profiling data (Explore1536) (19)
- BAFF-R expression level on B cells from malaria patients (26).
- iRBC stimulation of PBMCs from malaria patients (18).

All code used in the analyses is available via https://github.com/SundlingLab/MalariaMonocytes.

European Data Regulations, as interpreted by Karolinska Institutet, precludes open deposition of sensitive personal data into public repositories. This includes biological data that can be traced back to an individual, so-called pseudoanonymized data. The transcriptomic and proteomic data are available pending relevant permits via https://doi.org/10.48723/cmgs-aj72. This includes a metadata record describing the existing datasets. Data points in graphs derived from pseudoanonymized donors are available from the corresponding author, pending relevant permits. Values for all data points in graphs coming from anonymized donors or aggregated data are reported in the Supporting Data Values.

### Experimental model and subject details

#### Sex as a biological variable

The study examined immune responses during and after malaria with both sexes included, except for the single-cell RNA sequencing data, which was based on males only. This was done to reduce variability as the dataset is limited to four donors.

#### Study design/cohort

This study is based on new and repurposed data, previously generated on the peripheral blood of patients treated for *P. falciparum* malaria at Karolinska University Hospital. The cohort is well characterized and has been extensively described in the context of research questions related to antibody longevity in absence of re-exposure (61), B cell profiling (26), and antibody interplay (18), identification of antibody markers for recent malaria exposure (62), and protein-centric multi-omics blood profiling (19).

The prospective study was conducted at the Karolinska University Hospital in Stockholm, Sweden, with ethical approval from the Ethical Review Board in Stockholm and the Swedish Ethical Review Authority (Dnr 2006/893-31/4, 2013/550-32, 2015/2200-32, 2016/1940/32, 2018/2354-32, 2019-03436, 2020-00859, 2020-04147, 2024-00374-02). The work with PBMCs from healthy human blood was approved under permit Dnr 2025-01197-01. All participants gave written informed consent.

## Method details

### Bioinformatic and statistical analyses

All data wrangling and analysis were performed in R (v.4.3.2) using packages from the tidyverse (v.2.0.0) (63). If not stated otherwise, data was visualized using R packages ggplot2 (v.3.4.2), ComplexHeatmap (v.2.16.0), circlerize (v.0.4.15). The figures were assembled using patchwork (v.1.1.2) or Adobe Illustrator.

### Single-cell multi-omics data set

We used the previously reported targeted single-cell multi-omics data, generated on PBMCs from malaria-infected individuals within the Karolinska cohort (19). In short, CD45^+^ PBMCs from 4 individuals (2 primary infected, 2 previously exposed) with three-time points (Acute, D10, M12) were profiled on a transcriptional level using the Immune Response Panel (targeting 399 genes), as well as on a cell surface level using 29 AbSeq oligo-conjugated antibodies (BD Biosciences). A total of 73,121 cells were annotated based on the expression of 374 genes and 29 surface markers, resulting in 27 immune cell subsets as previously described (19). Here, we focused mainly on myeloid cells, annotated as pDCs, mDCs, CD14^+^ and CD16^+^ monocytes.

All downstream analyses were performed using the Seurat R package (v4) (64). To address interindividual variability and reduce biases associated with single-cell–level differential expression analyses, we adopted a pseudobulk approach. For each donor and time point, raw counts from all selected cells were aggregated using Seurat’s AggregateExpression() function to generate donor-level pseudobulk profiles. Aggregated counts were subsequently analyzed with DESeq2 (v1.40) through Seurat’s FindMarkers() function to identify genes differentially expressed between acute infection and later time points. Donor identity was included as a blocking factor in the design formula to account for subject-specific effects. Genes with a P value < 0.01 were considered statistically significant.

### Differential protein abundance analysis

Plasma protein data previously published by Lautenbach et al. (19) comprised longitudinal measurements collected during acute *Plasmodium falciparum* malaria and throughout convalescence in the same cohort of returning travelers. These data were used to assess changes in protein abundance across groups and sampling time points. Differential protein abundance analysis was performed using linear mixed-effects models implemented in the *lmerTest* package in R (65, 66), with time point included as a fixed effect and subject as a random effect to account for repeated measurements within individuals:

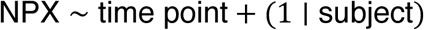

Post hoc pairwise comparisons were performed using the *emmeans* package (67) to identify significant differences both between sampling time points and between exposure groups at the acute time point. For group comparisons, the contrast *Acute primary_infected – Acute previously_exposed* was evaluated. Resulting *P* values were adjusted for multiple testing using the Benjamini–Hochberg procedure to control the false discovery rate(68). Statistical significance in figures is denoted as follows: *P* ≤ 0.05 (\**), P < 0.01 (**), P < 0.001 (\*\*\**), and *P* < 0.0001 (****).

### Protein annotation

Proteins were annotated using the Human Protein Atlas, version 24.0, retrieved from https://swww.proteinatlas.org/. DAPs were mapped based on “Secretome location” and “Secretome function” annotations.

### Stimulation of PBMC with iRBC

Flow cytometry data from a previously generated functional experiment was re-analyzed. The experiment was designed to assess functional differences in γ8 T cells in PBMCs from malaria-infected individuals (18). We repurposed the data and gated on CD14^+^ monocytes investigating specifically the stimulation with uninfected and infected RBCs and with PMA (phorbol 12-myristate 13-acetate) and ionomycin. See the gating strategy in **Figure S2**.

### Plasma pooling and antibody depletion

Plasmas were pooled from eight donors with primary infection or eight donors with previous malaria exposure with matched samples at the acute and 12-month time-points. The pools were depleted of IgG antibodies using protein-G spin columns (Ab SpinTrap, Cytiva) as described previously (23). IgG depletion was confirmed using an in-house multiplex bead assay detecting total IgG. Briefly, High-binding 96-well plates (Costar) were coated overnight at 4°C with goat anti-human IgG (Merck, I2136; 1 μg/mL in PBS, 100 μL/well). Plates were washed 4 times with PBS containing 0.05% Tween-20 (PBST) and blocked with 1% BSA and 0.05% Tween-20 in PBS (pH 7.4; 100 μL/well) for 1 hour at room temperature (RT). Plasma pools (pre-depletion, post-depletion, and eluted IgG fractions) were serially diluted starting at 1:100, followed by 5-fold serial dilutions (8 total), and added in duplicate (100 μL/well). Plates were incubated for 30 minutes at room temperature. After washing, peroxidase-conjugated goat anti-human IgG (Jackson ImmunoResearch; 1:10,000 in blocking buffer) was added (100 μL/well) and incubated for 30 minutes at RT. Plates were washed four times with PBST, and TMB substrate (Thermo Scientific; 75 μL/well) was added for 5-10 minutes before stopping the reaction with 0.2 M H₂SO₄ (75 μL/well). Absorbance was measured at 450 nm using a microplate reader (Thermo Scientific). Depletion was confirmed by comparing EC50 values, to range from 90.5-99.1% (**Figure S9**).

### Sorting of B cells

For the in vitro cultures of B cells, the cells were sorted from buffy coats from healthy donors. Buffy coats were obtained from the Karolinska University Hospital blood bank, in Stockholm, Sweden. Complete media consisted of RPMI-medium supplemented with 10% heat-inactivated FBS, 2mM L-glutamine, 100 U/ml of penicillin, 100 mg/mL of streptomycin, 10 mM HEPES and 0.05 mM 2-Mercaptoethanol (all from ThermoFisher Scientific). B cells were isolated through negative selection using either the RosetteSep Human B Cell Enrichment Cocktail (Stemcell technologies) as previously described(69) or with MACS cell separation (Milteny Biotech). Briefly, 25 μL of the Cell Enrichment Cocktail was added for each ml of blood, followed by incubation for 20 minutes at room-temperature (RT). Then two volumes of PBS + 2% FBS were added to the blood. 15 mL Ficoll–Paque Plus (GE Healthcare) was added to SepMate (Stemcell technologies) tubes and centrifuged at 1000 *g* for 1 min at room temperature (RT). The blood was poured into the SepMate tubes and centrifuged at 1200 *g* for 10 min with the brake on. The cell fraction was transferred to new tubes and washed twice with PBS + 2% FBS (300 *g*, 10 min, brake at 5 out of 9, RT). For the MACS cell separation, the B cells obtained from the RosettSep isolation were incubated with a Biotin-Antibody cocktail (Milteny Biotec) targeting non-B cells for 5 min at 4 °C, and then for another 10 min (4 °C) with the anti-Biotin Microbeads (Milteny Biotec). The cell suspension was then passed through LS columns placed in the MACS separator (Milteny Biotec). The B cell fraction was saved and washed as previously described. After the final wash, the isolated cells were resuspended in PBS + 2% FBS and counted using a Countess II (Invitrogen) following a 1:1 dilution with 0.4% trypan blue. The cells were centrifuged again and resuspended at 5 × 10^6^ cells/mL in freezing medium (FBS + 10% DMSO) and placed inside a CoolCell at – 80°C overnight, after which the cells were transferred to liquid nitrogen for long-term storage. B cell purity was assessed by flow cytometry and were on average±SD 92.3±6.6%.

### In vitro stimulation of sorted B cells and fresh PBMCs

For the in vitro stimulation, previously sorted B cells were thawed, moved to a 15 mL Falcon tube and 13 mL of complete media was added dropwise, the cell suspension was carefully mixed, and rested on ice for 20 min. The sorted B cells were then centrifuged at 300 *g* for 5 min, washed once in complete media (300 *g*, 5 min), resuspended in 1 mL complete media and counted, as previously described. Stimulations of purified B cells were done in a 96-well round-bottom tissue-culture plate (Techno Plastic Products). The stimuli included α-Ig (IgA+IgG+IgM [H+L] AffiniPureF(ab’)2 Fragment goat-anti-human [Jackson ImmunoResearch Laboratories]) at 1.25 µg/mL, CpG-C (Invivogen) at 2.5 µg/mL, and Human BAFF (PeproTech) at 125 ng/mL. The stimuli were added to wells with 1 × 10^5^ cells in a total volume of 200 µL complete media.

Fresh PBMCs were isolated from healthy donors and stimulated with pooled plasma from malaria patients. Plasma samples were collected either during acute infection (8 primary-infected and 8 previously exposed donors) or at 12 months post-treatment (donor matched 8 primary-infected and 8 previously exposed donors), with equal volume contributions from each donor. The total IgG from the pooled plasma have been removed by Ab SpinTrap (Cytiva). Freshly isolated PBMCs were cultured at 37°C and 5% CO_2_ for 20–24 hours in RPMI-1640 medium supplemented with either 20% IgG-depleted plasma (acute or convalescent, from either primary-infected or previously exposed donors) or 20% FBS. Plates were incubated at 37 °C and 5% CO_2_.

### Flow cytometry of stimulated B cells and stimulated fresh PBMCs

Cells were washed twice with PBS and stained with a Live/Dead viability dye (Thermo Fisher Scientific) for 20 minutes at 4°C. After washing twice with PBS containing 2% FBS, cells were incubated with an antibody cocktail for surface marker staining for 20 minutes at 4°C. For intracellular staining, cells were washed once after surface staining and fixed/permeabilized using FoxP3 Fix/Perm buffer (eBioscience) for 30 minutes at room temperature. Cells were then washed with FoxP3 Wash/Perm buffer and incubated with intracellular antibodies for 30 minutes at room temperature. Following staining, cells were washed with Wash/Perm buffer and resuspended in PBS containing 2% FBS. Stimulated B cells were acquired on a 5-laser BD LSRFortessa or a 4-laser Cytek Aurora (Cytek Biosciences) while stimulated fresh PBMCs were analyzed on a 5-laser Cytek Aurora (Cytek Biosciences) with CountBright absolute counting beads (Thermo Fisher Scientific). Analysis of flow cytometry data was done with FlowJo (version 10.10.0, BD for stimulated B cells and version 10.10.1, BD for stimulated fresh PBMCs). See **Table S2** for antibody panels.

### Multiplex bead-based assay to measure parasite-specific antibodies

A multiplex bead-based antibody assay was performed to measure total IgG, IgG1, IgG3, and IgM responses against recombinant merozoite surface proteins (AMA1, MSP1, MSP2, MSP3, GAMA derived from sequences of the *P. falciparum* 3D7 parasite strain). The plasma samples used for this study were collected from participants at the day of diagnosis when they visited the Karolinska University hospital. Briefly, the plasma samples were diluted at 1:250 with dilution buffer (0.1% BSA, 0.05% Tween-20 in PBS, pH 7.4). Related merozoites antigens were coupled to magnetic COOH beads on different regions (B27, 35, 44, 52, 62, Bio-Rad). Prior to antigen coupling, buffer exchange was done to phosphate-buffered saline (PBS) using Zeba spin desalting columns (Thermo Scientific). Antigen and magnetic beads coupling were performed with the Bio-Plex Amine coupling kit (Bio-Rad), EDAC (Bio-Rad) and Sulfo-NHS (Thermo Scientific). The antigen-coupled beads were diluted with dilution buffer to 50 μl, corresponding to 1250 beads per region and mixed with 50 μl 1:250 diluted plasma in 96-well microtiter plates (Bio-Rad), followed by 1 hour incubation on a shaker at room temperature avoiding light. After washing 4 times with PBS+0.05% Tween-20, 100 μl of PE-conjugated Goat Anti-Human IgG (Jackson ImmunoResearch Laboratories) diluted at 1:600 or Hinge PE mouse-anti-human IgG1 (diluted at 1:300), IgG3 (diluted at 1:300), or IgM (diluted 1:300) (all from Southern Biotech) was added for a 30 min incubation on a shaker at room temperature, followed by washing another 4 times with PBS+0.05% Tween-20. The beads were then resuspended with 100 μl dilution buffer for sample acquisition. Median fluorescent intensities (MFIs) were measured using the Bio-Plex 200 system (Bio-Rad), with a reader setting to read a minimum of 100 beads per bead region.

### Statistical analyses

Statistical analyses were performed using GraphPad prism version 11 or R and are further described in the figure legends. Briefly, analyses comparing donor responses or characteristics over time or following multiple stimulations were done using repeated measures two-tailed one-way or two-way ANOVA. If the analyses included missing data, a linear mixed effects model was used. For comparison between two variables, two-tailed student’s t-tests or Wilcoxon rank tests were used. P-values < 0.05 were considered significant with \**P* < 0.05, ** *P* < 0.01, \*\*\**P* < 0.001, and \*\*\*\**P* < 0.0001.

## Additional information

## Acknowledgement

We would like to thank all study participants for contributing to the study with their blood and time. Moreover, we would also like to express our gratitude to all clinicians, nurses, laboratory staff, and students for assisting us with study inclusion, sampling, and processing. We thank all colleagues and peers for valuable feedback during the process of this study. We gratefully acknowledge the Human Protein Atlas for providing invaluable resources, with special thanks to Mathias Uhlén and Fredrik Edfors for their leadership in the Disease Atlas initiative, generously supported by the Knut and Alice Wallenberg Foundation. We acknowledge support from the National Genomics Infrastructure in Stockholm funded by Science for Life Laboratory, the Knut and Alice Wallenberg Foundation and the Swedish Research Council, and SNIC/Uppsala Multidisciplinary Center for Advanced Computational Science for assistance with massively parallel sequencing and access to the UPPMAX computational infrastructure.

## Funding support

The study was supported by the following grants:

- From Vetenskapsrådet to CS: 2019-0194 and 2023-01943
- From the Åke Wiberg foundation to CS: M18-0076.
- From Magnus Bergvall foundation to CS: 2017-02043 and 2018-02656
- From Karolinska Institutet to CS: FS-2020:0007 and FS-2022:0010
- From Karolinska Institutet to AF: 2019-00992.
- From Vetenskapsrådet to AF for the cohort: 521-2012-3311, 2015-02977, 2018-02688, 2018-04468, 2021-03105
- From Region Stockholm to AF for the cohort: 20150135, 20180409, 960104, 986923

## Author contribution

M.J.L. and C.S. designed the study. A.F. was responsible for the cohort. L.K., P.X., C.SS., A-D.CG., F.C., and M.S.G. performed functional experiments. M.J.L. performed bioinformatic analysis. All authors interpreted the results. M.J.L., P.X., and C.S. generated figures. C.S. supervised the study. A.F. and C.S. provided funding. M.J.L. wrote the original draft with input from C.S. M.J.L, P.X., and C.S. revised the study. All authors contributed to editing.

## Declaration of generative AI and AI-assisted technologies in the writing process

During the preparation of this work, the authors used Microsoft’s Copilot to improve wording and clarity. After using this tool/service, the authors reviewed and edited the content as needed and take full responsibility for the content of the publication.

