## Supplemental material for "Previous malaria exposure attenuates monocyte-driven inflammation and correlates with modulation of the B cell response"

### Supplementary Material

#### Table S1. Demographics

| Characteristic | Statistic | Number of participants | Primary infected | Previously exposed | Healthy control | P-value |
|---|---|---|---|---|---|---|
| **Related to Figure 2D-E, and Figure 4** | | | | | | |
| **Age, (years)** | Median (Range) | 72 | 34 (20, 60) | 41 (27, 63) | - | 0.060 |
| **Time since last residency in endemic area, (years)** | Median (Range) | 41 | - | 12 (0, 33) | - | |
| **Cumulative time of residency in endemic area, (years)** | Median (Range) | 44 | - | 26 (4, 41) | - | |
| **Female sex** | n (%) | 72 | 6 (25%) | 11 (23%) | - | >0.9 |
| **CMV positive** | n (%) | 67 | 16 (70%) | 43 (98%) | - | 0.002 |
| **Related to Figure 3A-B** | | | | | | |
| **Age, (years)** | Median (Range) | 11 | 38 (20, 53) | 49 (27, 63) | - | 0.8 |
| **Time since last residency in endemic area, (years)** | Median (Range) | 6 | - | 16 (1, 32) | | |
| **Cumulative time of residency in endemic area, (years)** | Median (Range) | 6 | - | 24 (18, 39) | | |
| **Female sex** | n (%) | 11 | 1 (25%) | 0 (0%) | - | 0.5 |
| **CMV positive** | n (%) | 11 | 3 (60%) | 6 (100%) | - | 0.2 |
| **Related to Figure 5** | | | | | | |
| **Age, (years)** | Median (Range) | 57 | 34 (20, 60) | 37 (27, 63) | 32 (21, 57) | 0.8 |
| **Time since last residency in endemic area, (years)** | Median (Range) | 28 | - | 10 (0, 32) | - | |
| **Cumulative time of residency in endemic area, (years)** | Median (Range) | 28 | - | 26 (15, 39) | - | |
| **Female sex** | n (%) | 57 | 4 (27%) | 4 (14%) | 10 (71%) | 0.4 |
| **CMV positive** | n (%) | 43 | 10 (67%) | 27 (96%) | - | 0.015 |
| **Related to Figure 3 and Figure 6 plasma pools** | | | | | | |

| | | | | | | |
|---|---|---|---|---|---|---|
| **Age, (years)** | Median (Range) | 16 | 41 (27, 53) | 45 (27, 63) | - | >0.9 |
| **Time since last residency in endemic area, (years)** | Median (Range) | 8 | | 14 (1, 32) | | |
| **Cumulative time of residency in endemic area, (years)** | Median (Range) | 8 | - | 25 (18,39) | - | |
| **Female sex** | n (%) | 16 | 4 (50%) | 1 (13%) | - | 0.3 |
| **CMV positive** | n (%) | 16 | 3 (38%) | 8 (100%) | - | 0.026 |
| **Related to Figure 7** | | | | | | |
| **Age, (years)** | Median (Range) | 55 | 35 (20, 60) | 40 (27, 57) | - | 0.3 |
| **Time since last residency in endemic area, (years)** | Median (Range) | 30 | - | 13 (0, 33) | - | |
| **Cumulative time of residency in endemic area, (years)** | Median (Range) | 33 | - | 26 (10, 39) | - | |
| **Female sex** | n (%) | 55 | 6 (30%) | 8 (23%) | - | 0.7 |
| **CMV positive** | n (%) | 55 | 14 (70%) | 34 (97%) | - | 0.007 |

*P-values compare primary infected and previously exposed donors only. Continuous variables were analyzed using the Wilcoxon rank-sum test; categorical variables were analyzed using Fisher's exact test. Healthy controls were excluded from statistical comparisons.*

#### Table S2. Flow cytometry panels

| Antibodies used for plasma stimulation flow panel | | | |
|---|---|---|---|
| **Antibodies** | **Clone** | **Source** | **Identifier** |
| CD56 BB700 | NCAM16.2 | BD | Cat#566573, RRID:AB_2744430 |
| IgM BB515 | G20-127 | BD | Cat#564622, RRID:AB_2738869 |
| CD69 PE-Cy7 | FN50 | BD | Cat#557745, RRID:AB_396851 |
| CD86 PE-Cy5 | IT2.2 | BioLegend | Cat#305408, RRID:AB_314528 |
| CD307e (FcRL5) PE | 509f6 | BioLegend | Cat#340304, RRID:AB_2104588 |
| CD3 APC-H7 | SK7 | BD | Cat#560176, RRID:AB_1645475 |
| CD16 R718 | B73.1 | BD | Cat#567228, RRID:AB_3683755 |
| CD268 (BAFF receptor) AF647 | 11C1 | BD | Cat#564817, RRID:AB_2744347 |
| CD11c BV786 | B-ly6 | BD | Cat#568220, RRID:AB_2916848 |
| CD14 BV711 | MψP9 | BD | Cat#563372, RRID:AB_2744290 |
| CD27 BV650 | M-T271 | BD | Cat#564894, RRID:AB_2739004 |
| HLA-DR BV510 | G46-6 | BD | Cat#563083, RRID:AB_2737994 |
| CD64 BV421 | 10.1 | BD | Cat#562872, RRID:AB_2737856 |
| IgD BUV737 | IA6-2 | BD | Cat#612798, RRID:AB_2870125 |
| CD19 BUV496 | SJ25C1 | BD | Cat#612938, RRID:AB_2870221 |
| Green fluorescent reactive dye | | Invitrogen | Cat#L34970 A |
| CountBright absolute counting beads | | Thermo Scientific | Cat#C36950 |

| Antibodies used for B cell stimulation flow panel | | | |
|---|---|---|---|
| **Antibodies** | **Clone** | **Source** | **Identifier** |
| Aqua fluorescent reactive dye | | Invitrogen | L34957 |
| CD38 BV421 | HIT2 | BD | Cat#562444, RRID:AB_11151894 |
| CD27 BV650 | M-T271 | BD | Cat#564894, RRID:AB_2739004 |
| CD3 PE-Cy5 | UCHT1 | BD | Cat#300410, RRID:AB_314064 |
| CD14 PE-Cy5 | M5E2 | BD | Cat#301864, RRID:AB_2860767 |
| CD19 PE-Cy7 | HIB19 | BD | Cat#560728, RRID:AB_1727438 |
| CD268 (BAFF receptor) AF647 | 11C1 | BD | Cat#564817, RRID:AB_2744347 |
| CD20 APC-H7 | 2H7 | BD | Cat#641405, RRID:AB_1645729 |

#### Figure S1

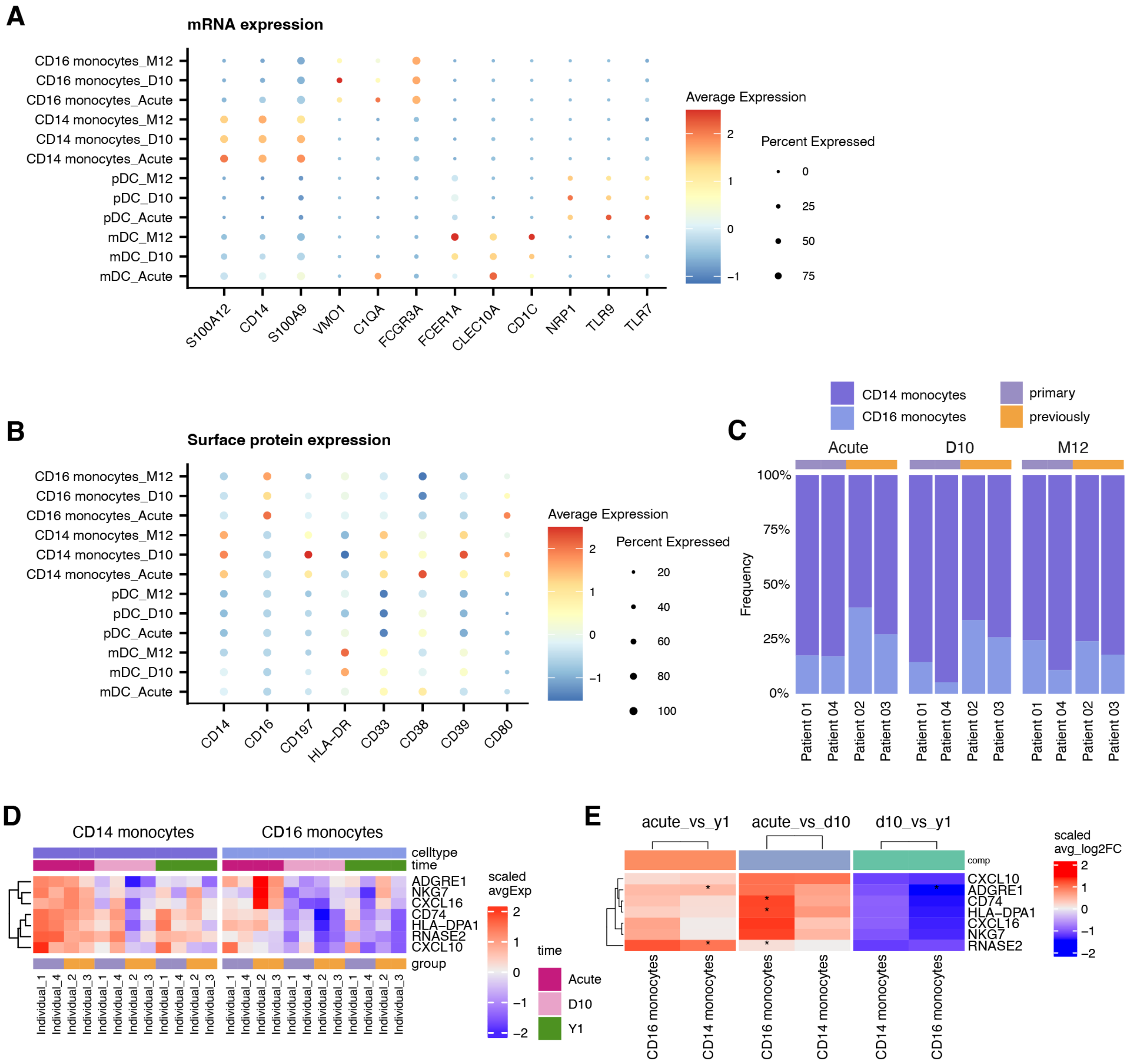


***Figure S1. Gene and protein expression of myeloid cell-specific markers, r*elated to main Figure 1.** *(**A**) Gene expression in myeloid cells. (**B**) Surface protein expression on myeloid cells. (**C**) Distribution of CD14 and CD16 monocytes for each donor at each time-point. (**D**) Pseudobulk expression of indicated genes for each donor in CD14 and CD16 monocytes. (**E**) Grouped pseudobulk expression of selected genes across monocyte subsets and time points. Statistical analysis for G and I was performed using DESeq2 on individual pseudobulk data with *P < 0.01.*

### Figure S2

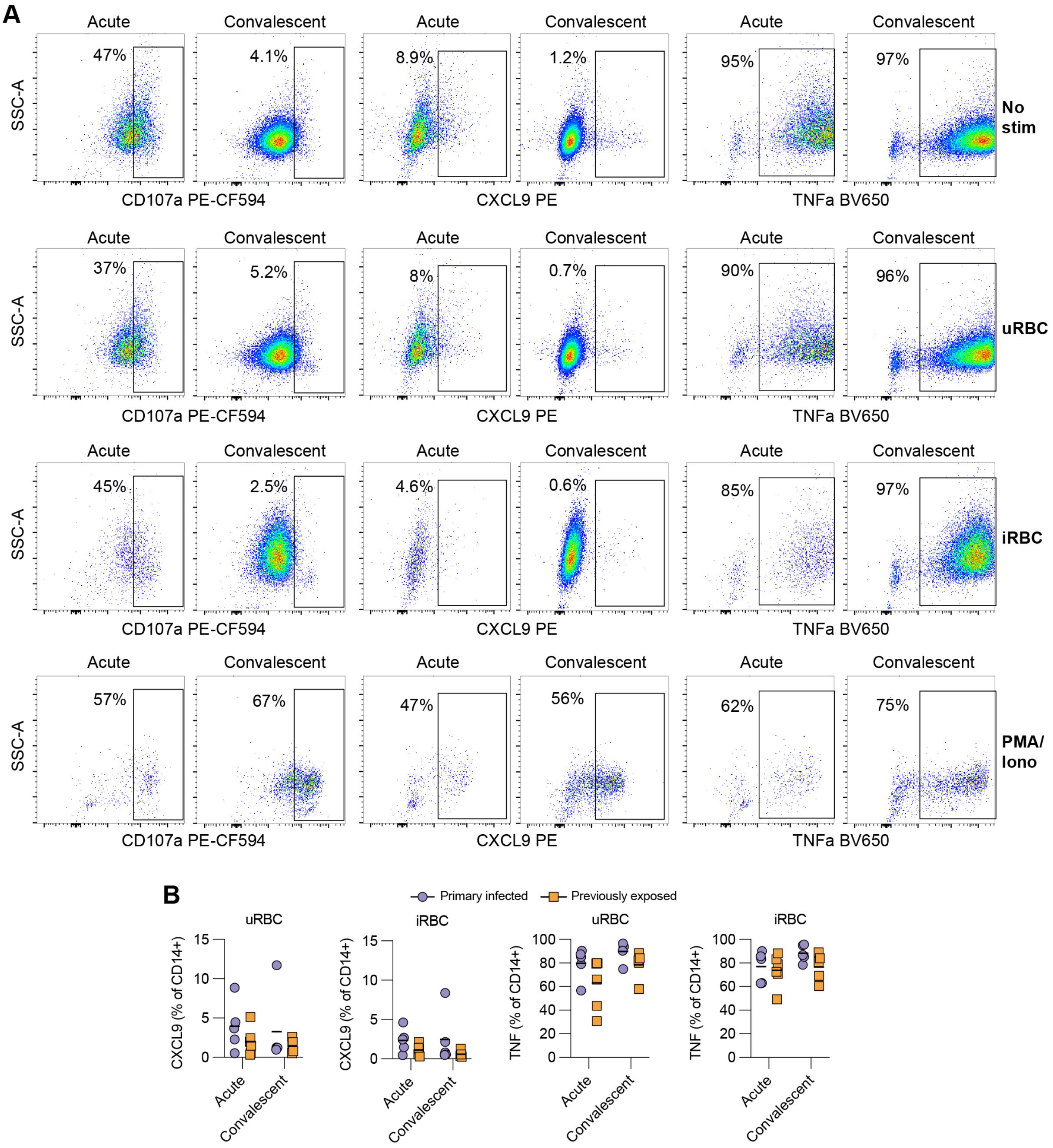


***Figure S2. Gating strategy to assess monocyte CD107a, CXCL9, and TNF-alpha in stimulated PBMCs, related to main Figure 3. (A)*** *Representative gating strategy to assess the frequency of CD107a, CXCL9, or TNFa-positive Live, CD3–CD57–CD14+ monocytes. Stimulation is indicated on the right with cells receiving no stimulation (media), uninfected RBC (uRBC), infected RBC (iRBC), or phorbol 12-myristate 13-acetate (PMA) and ionomycin (PMA/Iono) stimulation. (**B**) Frequency of CXCL9 and TNFa-positive cells out of total CD14+ monocytes in stimulations with uRBC or iRBC. Statistical analysis was done using a linear mixed effects model followed by Sidak's posttest for correction for multiple testing. Statistical analysis was done using a matched pair two-tailed two-way ANOVA followed by Sidak's posttest. P < 0.05 were considered significant. Only significant p-values are shown.*

#### Figure S3

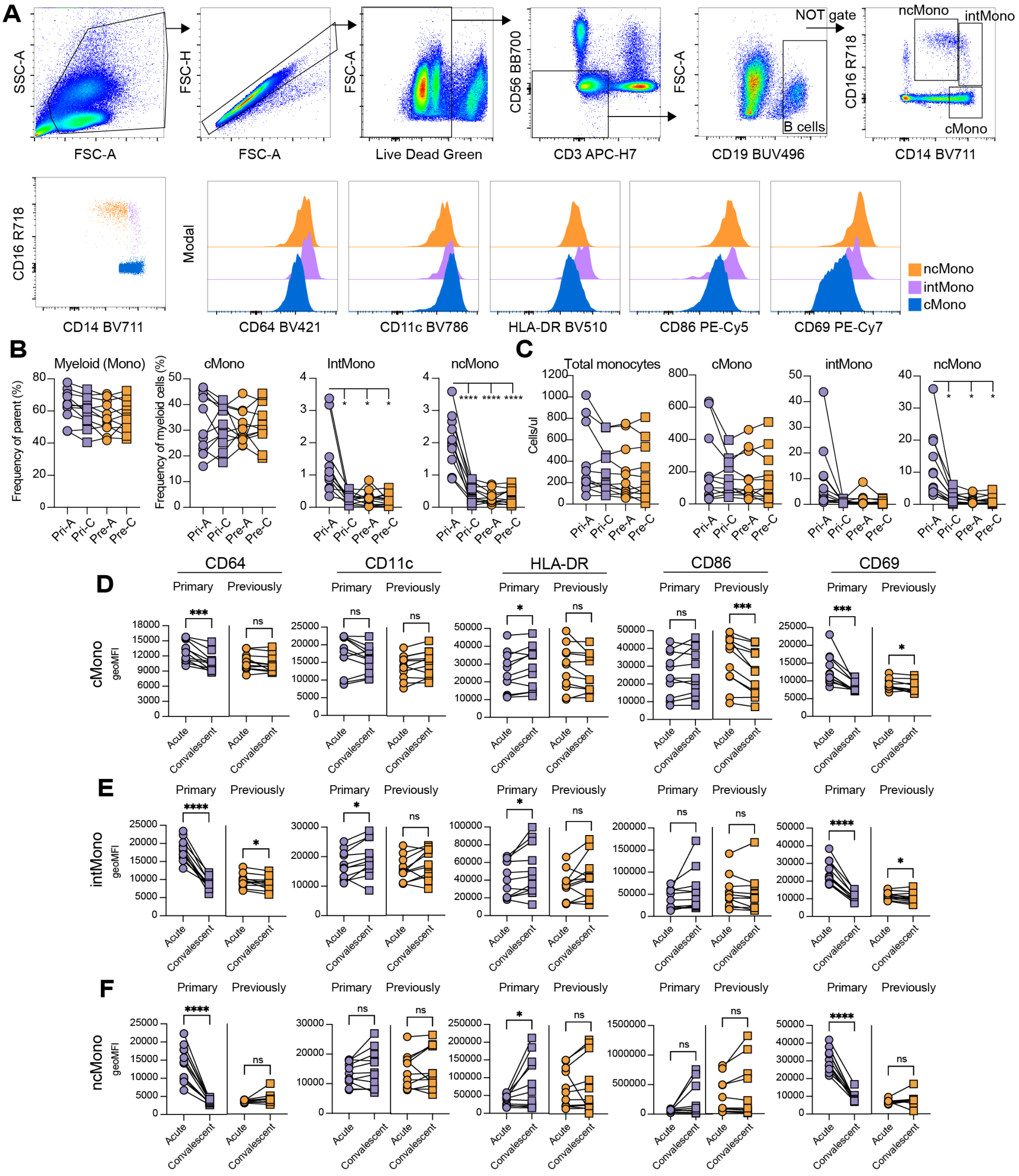


*Figure S3. Impact of patient plasma stimulation on monocytes, related to main Figure 3.* (***A***) *Gating strategy for monocytes.* (***B***) *Frequency of parent (live, non-B/T/NK cells).* (***C***) *Cell counts of total monocytes and monocyte subsets after 24 h PBMC culture with patient plasma pools from primary infected individuals (pri, purple) and previously exposed individuals (pre, orange) at acute malaria (A, circles) and 12 months post-infection (C, convalescent, boxes).* (***E-F***) *Geometric mean fluorescent intensities (gMFI) for CD64, CD11c, HLA-DR, CD86, and CD69 on* (***C***) *classical monocytes,* (***D***) *intermediate monocytes, and* (***E***) *non-classical monocytes. Statistical analyses for C and D were done using two-way matched pair ANOVA with Geisser-Greenhouse correction followed by Tukey's post-test. Analyses for D-F were done using a two-tailed paired student's t-test. Ns $p > 0.05$, *$p < 0.05$, **$p < 0.01$, ***$p < 0.001$, ****$p < 0.0001$.*

### Figure S4

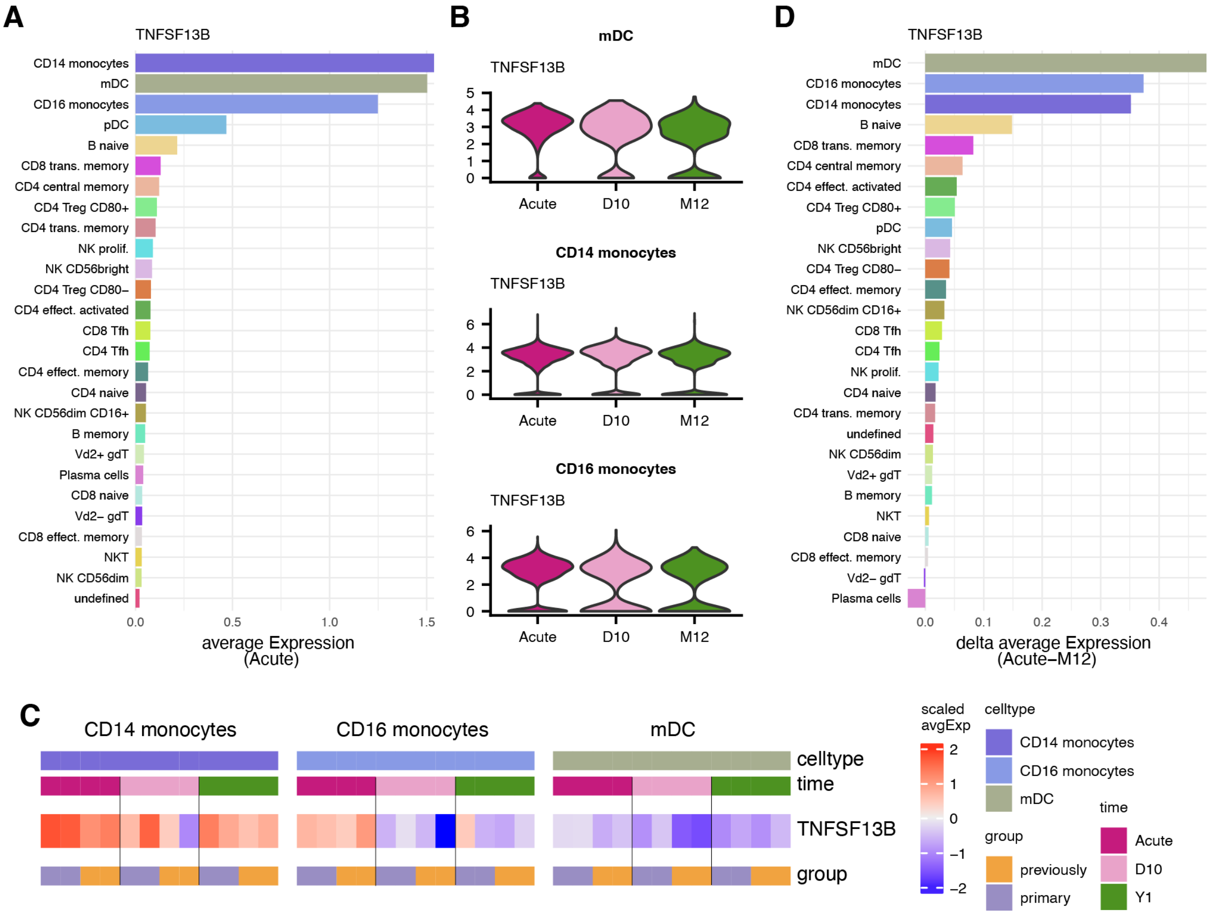


***Figure S4. Monocytes and dendritic cells in patients with malaria express the gene encoding for BAFF, related to main Figure 5.*** *(**A**) The average gene expression of BAFF encoding gene (TNFRSF13) per immune cell subset in PBMCs of four individuals with acute malaria. (**B**) Gene expression of TNFRSF13 over sampling time points, acute (pink), 10 days after (light pink) and 12 months later (green). (**C**) Individual-specific pseudobulk expression of TNFSF13B across myeloid cell subsets and time points. (**D**) Delta average gene expression (Acute-M12) of all immune cell subsets from four individuals.*

#### Figure S5

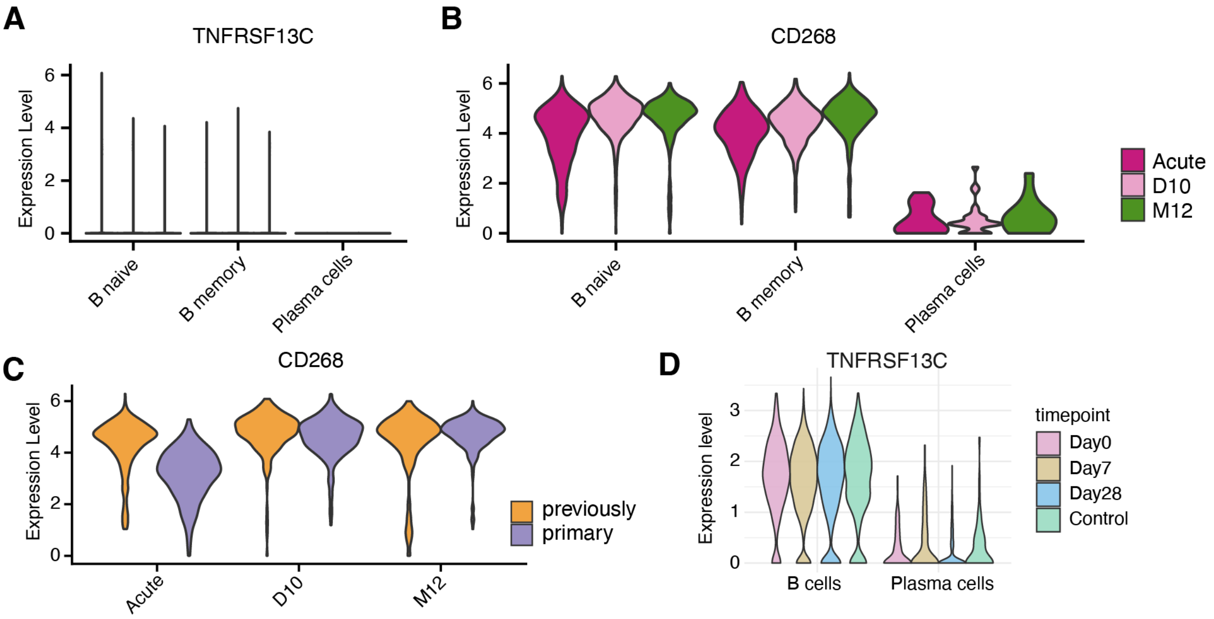


***Figure S5. Gene and protein expression on B cells, r*elated to main Figure 5.** *(**A**) Gene expression of TNFRSF13C (BAFF-R) on B cells. (B) CD268 (BAFF-R) protein levels on the cell surface of B cells. (**C**) Differences in CD268/BAFF-R protein levels on naïve B cells during acute malaria and 10 days and 12 months after treatment. (**D**) Repurposed single cell RNA sequencing expression levels from Dooley et al. (25) of TNFRSF13C (BAFF-R) in malaria patients (n = 6 donors) at acute infection (pink), and 7 days (brown), and 28 days (blue) after treatment, and in controls (green, n=2 donors).*

### Figure S6

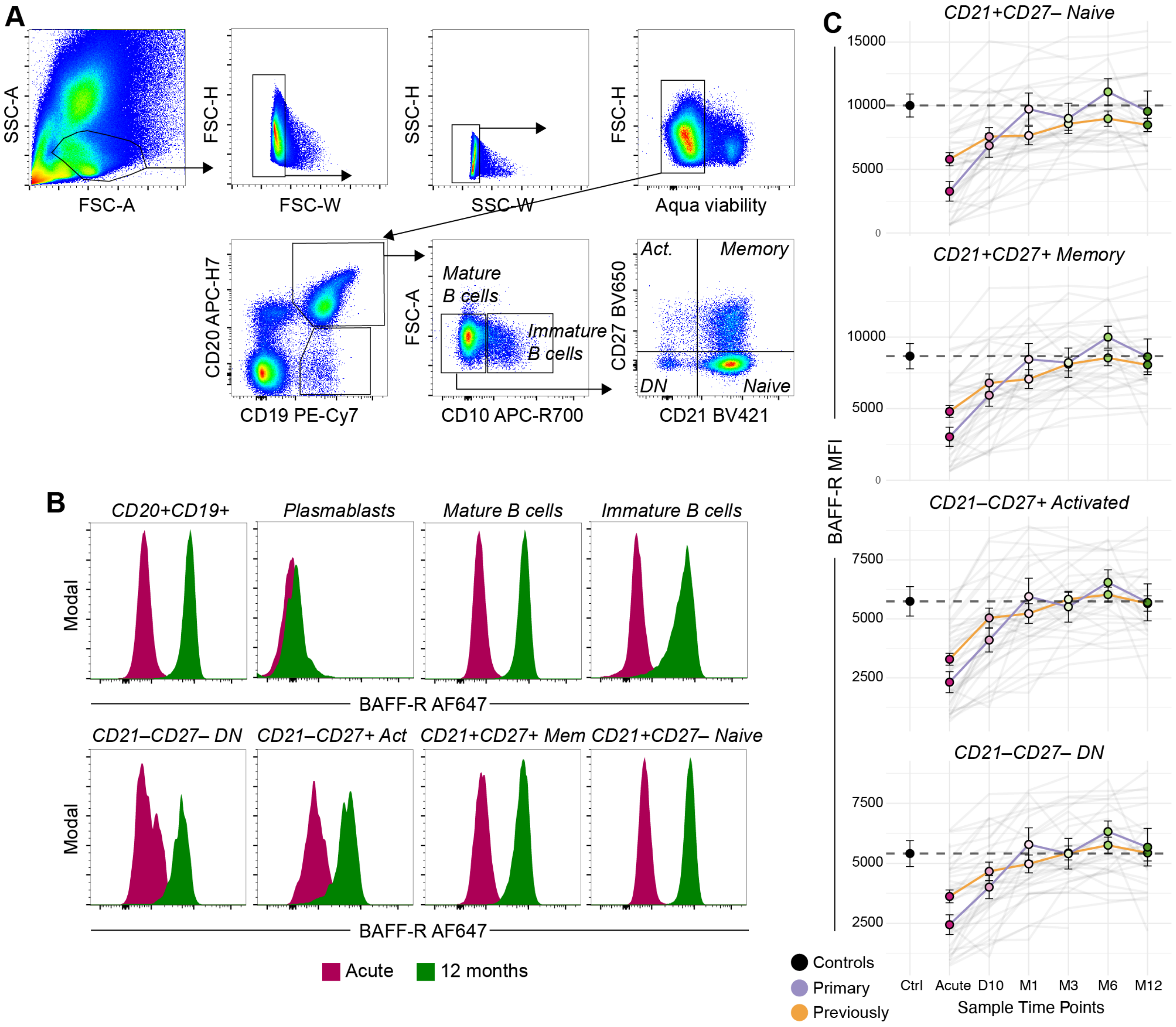


***Figure S6. BAFF-R gating and levels on B cell subsets, related to main Figure 5****. (****A****) Representative gating strategy for B cell subsets. (****B****) Overlay histograms of BAFF expression levels in different B cell subsets at Acute malaria (red) and 12 months after infection (green). (****C****) BAFF-R median fluorescent intensities of primary infected (purple, n=17) and previously exposed (orange, n=34) donors at different time-points following acute malaria for indicated B cell subsets. The mean BAFF-R of healthy controls (black, n=14) is indicated by a dashed line for each cell subset.*

### Figure S7

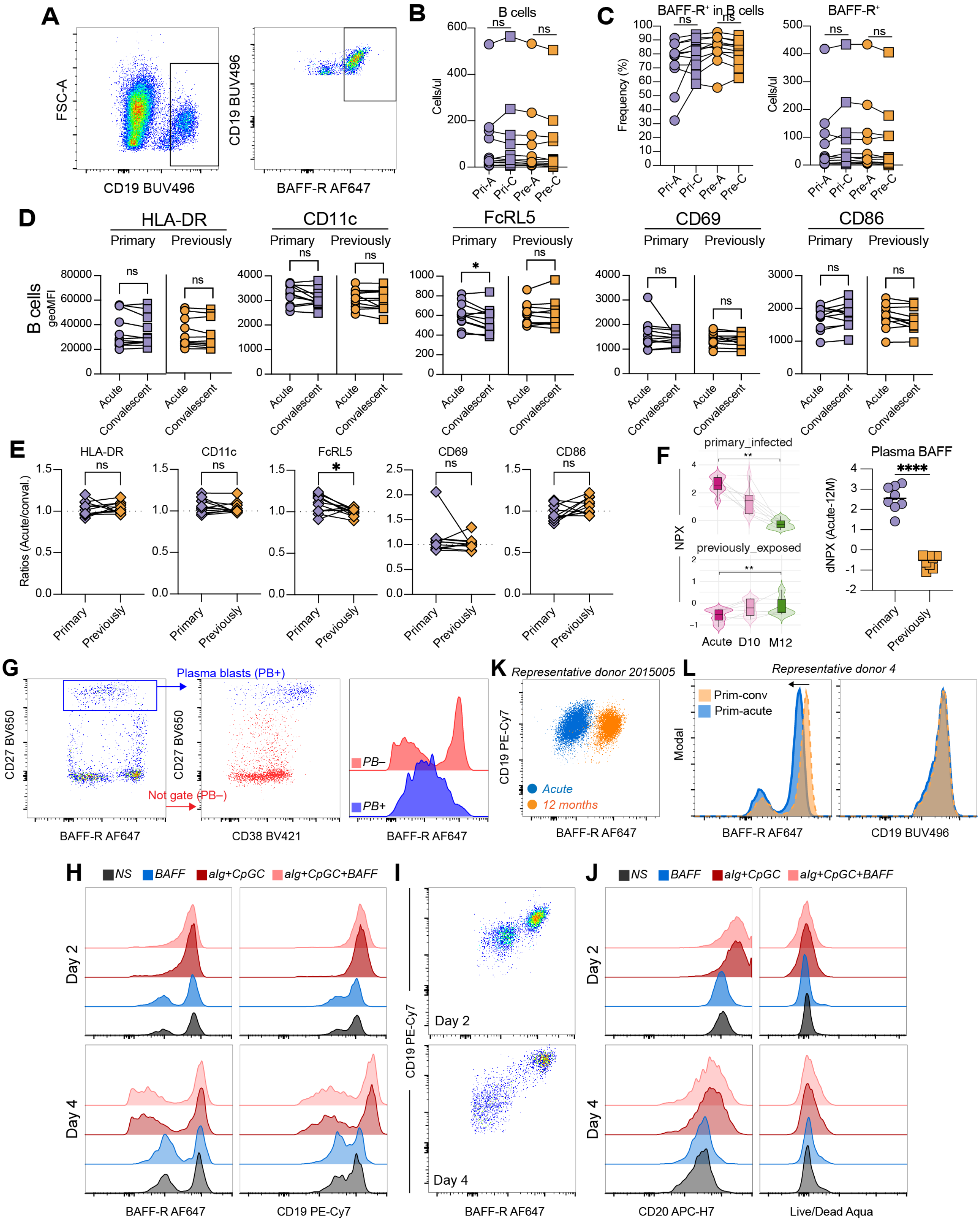


***Figure S7. BAFF-R dynamics following in vitro stimulation,*** **related to main Figure 6.** *(**A**) Representative gating for B cells followed by BAFF-R+ cells. (**B**) CD19+ B cells per microliter. (**C**) Frequency (left) and number (right) of BAFF-R+ cells of B cells. (**D**) Geometric mean fluorescent intensity (geoMFI) of indicated surface proteins on B cells after stimulation with indicated plasma pools for 24 h. (**E**) Ratios (acute/convalescent plasma) for geoMFI marker expression between cells stimulated with primary infected or previously exposed plasma pools. (**F**) BAFF*

*NPX (Acute, 10 days and 12 months) and delta-NPX (Acute–12 months) levels for donors included in the plasma pools. (**G**) BAFF-R versus CD27 at day 4 to gate for plasma blasts seen as CD27 high (PB+, blue) and non-PB (PB–, red) (left panel). Overlay of gated populations to indicate CD38 expression for PB+ and PB– cells (middle panel). Histograms indicating BAFF-R levels for PB+ and PB– cells. (**H**) Histograms indicating BAFF-R and CD19 levels on sorted stimulated B cells after 2 days (top) and 4 days (bottom) of stimulation with type of stimulation indicated by color. (**I**) Pseudocolor dot plots indicating CD19 and BAFF-R levels on stimulated non-PC B cells at day 2 and 4 of stimulation with BAFF alone. (**J**) Histograms indicating CD20 (left) and Live dead Aqua viability dye (right) levels at day 2 (top) and day 4 (bottom) of stimulation. (**K**) CD19 vs BAFF-R ex vivo levels from one representative donor with acute (blue) malaria and at 12 months post treatment (orange). (**L**) Histogram overlay of B cells gated from PBMCs stimulated with acute (orange) or convalescent (blue) plasma from primary infected donors. Arrow indicates shift in BAFF-R high levels. Statistical analyses in B-D was done using two-tailed paired student's t-tests while in E, F it was done using two-tailed unpaired student's t-tests with ns = p > 0.05, *p < 0.05, **p < 0.01, ***p < 0.001, ****p < 0.0001.*

#### Figure S8

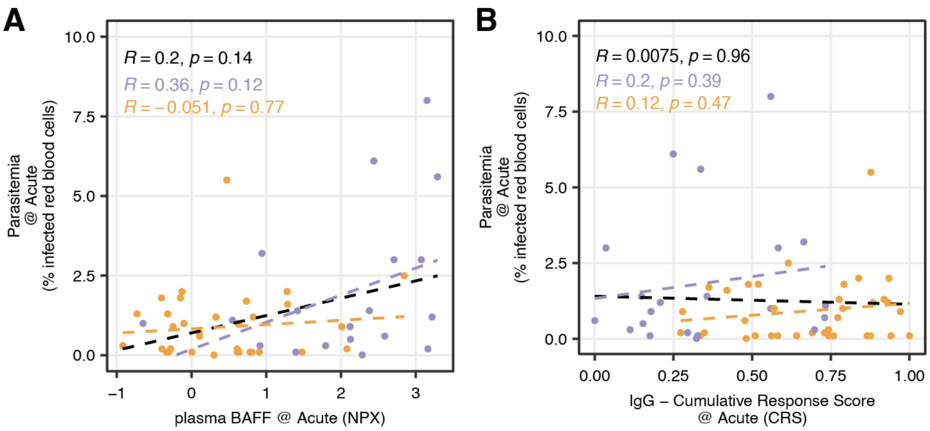


***Figure S8. Malaria parasitemia is not correlated with plasma BAFF levels or parasite-specific IgG, related to main Figure 7.*** *(**A**) Spearman-correlation of parasitemia with (**A**) plasma BAFF levels left and (**B**) cumulative antibody response score (CRS). Dots are colored by status of previous malaria exposure (orange n=35) or primary infection (purple, n=20).*

#### Figure S9

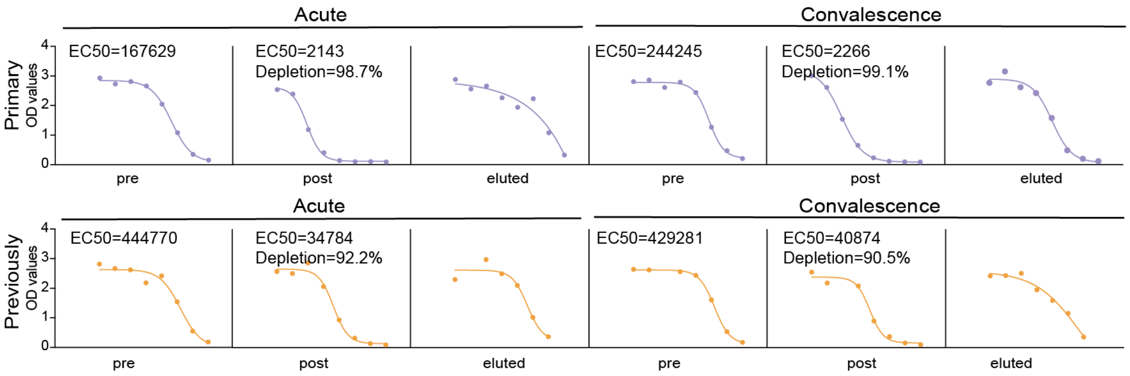


***Figure S9. Plasma pool antibody depletion**. Confirmation of antibody depletion in the four different plasma pools. Depletion was calculated as (1–(EC50-post / EC50-pre))*100.*
